# TGFβ-dependent upregulation of OCIAD2 is essential for epithelial-to-mesenchymal transition during mesendoderm differentiation

**DOI:** 10.1101/2025.11.10.687537

**Authors:** Kajal Kamat, Maneesha S. Inamdar

## Abstract

Deciphering mechanisms that govern lineage commitment of human pluripotent stem cells (hPSCs) is essential for optimizing differentiation strategies. Here, we investigated the role of the mitochondrial protein OCIAD2 in hPSC differentiation. We show that OCIAD2 is transiently upregulated during mesendoderm differentiation, in a TGFβ/Activin A-dependent manner. OCIAD2 depletion impairs mesendoderm induction and specification, resulting in an incomplete epithelial-to-mesenchymal transition (EMT). Through transcriptome analysis, immunoblotting and localization studies on OCIAD2-depleted (KO) or overexpressing (OV) human embryonic stem cells (hESCs), we identified OCIAD2 as a positive regulator of TGFβ signalling. Additionally, KO hESCs exhibited downregulated fatty acid oxidation (FAO) genes, indicating dysregulated lipid metabolism. Pharmacological restoration of FAO improved mesendoderm differentiation capacity in KO, suggesting a key role for OCIAD2 in coordinating metabolism and TGFβ signalling. We propose that OCIAD2 regulates EMT by integrating metabolic cues with TGFβ pathway activation. Our findings provide insight into how mitochondrial proteins regulate lineage commitment and EMT, with broader implications for both developmental biology and tumor progression.

**Highlights:**

- OCIAD2 expression peaks during mesendoderm specification of hESCs
- OCIAD2 depletion transcriptionally represses TGFβ signalling and impedes EMT
- Loss of OCIAD2 leads to hyperfused mitochondria and downregulation of FAO genes
- Pharmacologic supplementation with acetate enhances mesendoderm specification

## Introduction

Human pluripotent stem cells (hPSCs) possess the remarkable ability to self-renew indefinitely and generate all somatic cell types, offering unprecedented potential for regenerative medicine. Realizing this promise requires the generation and expansion of lineage-restricted progenitors *in vitro*, informed by developmental biology. During embryogenesis, a network of signalling pathways is repeatedly utilized to orchestrate tightly regulated transitions, resulting in diverse cell fates^1^. Despite advances in understanding external cues, the intrinsic molecular determinants of lineage commitment remain incompletely elucidated.

During vertebrate development, primitive streak (PS) formation from the pluripotent epiblast requires cells to ingress, undergo epithelial-to-mesenchymal transition (EMT), and migrate to generate the mesodermal and endodermal lineages. A coordinated interplay of BMP, WNT, Nodal, and FGF signalling, ensures spatial and temporal fidelity of development^2, 3^. A diverse array of transcription factors, RNA-binding proteins, and chromatin-related factors play key roles in establishing mesendoderm fate^4–7^. Additionally, mitochondrial bioenergetics and metabolic reprogramming also control mesendoderm differentiation^8–11^. Recent reports demonstrate that metabolic activity is not merely an outcome of cell state transitions, but actively regulates signalling pathways during murine gastrulation and axis segmentation, independent of its canonical role in energy generation^12–15^. However, molecular mechanisms by which metabolism interfaces with upstream signal activation and transduction are not well understood. To explore this, we studied the role of the mitochondrial protein Ovarian Carcinoma Immunoreactive Antigen Domain containing protein 2 (OCIAD2) in early lineage specification, using human embryonic stem cell (hESC) differentiation as a model.

The OCIAD protein family, comprising of OCIAD1 and OCIAD2, is implicated in several human pathologies, including carcinomas and neurodegenerative disorders^16–18^. The first identified member, OCIAD1/Asrij, is a stem cell-associated protein that maintains the hematopoietic stem cell pool in *Drosophila* and murine models^19–23^. While Asrij is essential for maintaining pluripotency of mouse ESCs^21^, OCIAD1 depletion from hESCs does not affect pluripotency. However, loss of OCIAD1 elevates mitochondrial Complex I activity and renders hESCs prone to differentiation^24^.

OCIAD2, the genomic neighbor of OCIAD1, arose by gene duplication in vertebrates^22^. OCIAD2 is expressed in multiple human fetal and adult tissues^16^ and is frequently detected in tumor samples, where its expression may be elevated or reduced, depending on the tumor type^25–27^. Functionally, OCIAD2 localizes predominantly to mitochondria and facilitates STAT3 activation and mitochondrial Complex III assembly in HEK293 cells^22, 28^. Its loss disrupts mitochondrial structure and promotes mitochondria-mediated apoptosis in lung adenocarcinoma cells^29^. Protein interaction network analysis of cancer datasets predicted that OCIAD2 is a TGFβ-responsive gene^30^. Further, clinical data from metastatic cancer patients support a positive correlation between OCIAD2 levels and TGFβ pathway activation^31^. Knockdown of OCIAD2 leads to increased AKT signalling, resulting in enhanced colony formation, migration, and invasion in hepatocellular carcinoma cells^27^. Recently, OCIAD2 was shown to promote glycolysis via AKT activation in pancreatic adenocarcinoma cells^26^.

A comparative single-cell transcriptomic analysis of human embryonic development reveals that OCIAD2 is primarily expressed in the epiblast, primitive streak, and mesendoderm lineages^32^. However, its physiological role, particularly in the context of early human development, remains unexplored.

We showed earlier that altering OCIAD2 levels does not affect pluripotency maintenance and tri-lineage differentiation capabilities of hESCs^33, 34^. In this study, through genetic manipulation, transcriptomic analysis and phenotypic characterization, we uncover a previously unreported functional link between OCIAD2-mediated metabolic regulation and TGFβ signalling, which impacts mesendoderm differentiation. Our findings may be leveraged to enhance stem cell differentiation efficiency and improve outcomes in directed differentiation protocols.

## Results

### OCIAD2 is upregulated at early stages of mesendoderm induction

Bulk transcriptomic (StemCellDB: https://stemcelldb.nih.gov/geneInformation/12538/I) and proteomic datasets^35^ generated from several human pluripotent stem cell lines report OCIAD2 expression in hESCs. By transcript and protein expression analysis of hESC line BJNhem20^36, 37^, we confirmed that undifferentiated hESCs express OCIAD2 (Figure S1A-C). Mining an integrated single-cell transcriptomic dataset of the zygote to gastrula stages of human embryogenesis^32^ indicated that *OCIAD2* expression levels are highest in the epiblast, PS and emerging mesendoderm lineages of the embryo (Figure S1D). In agreement with this, enrichment of *OCIAD2* transcript was seen in the transcriptome of undifferentiated hESCs and the emerging PS-like populations (Figure S1E), derived from directed mesoderm differentiation of hESCs^38, 39^. These data suggest a possible role for OCIAD2 in pluripotency and/or in mesendoderm specification.

For a finer mapping of OCIAD2 expression in hESC differentiation, we subjected BJNhem20 cells to a directed differentiation protocol (Figure S2A) that generates mesendoderm progenitors between day 3.5 to day 4.5^24, 40^. *OCIAD2* transcript level decreased as differentiation progressed (Figure S2B), consistent with the gene expression analysis reported previously^40^. However, OCIAD2 protein levels increased 1.5-fold during the initial days of differentiation (up to day 2.5), before declining sharply by day 4.5 (Figure 1A). This suggests complex regulation of OCIAD2 expression at early stages of differentiation. To evaluate the immediate effects of differentiation cues on OCIAD2 expression, we assessed protein levels in the first few hours of the onset of differentiation (at 0, 3, 6, 12 and 24 hrs post-induction). Expectedly, there was a marked increase (1.5-fold) in OCIAD2 protein levels within 3 hrs of differentiation (Figure 1B). This indicates that hESCs upregulate OCIAD2 upon mesendoderm induction, suggesting that it may be required for initiating differentiation. Since the mesendoderm differentiation protocol depends on direct activation of BMP, TGFβ and FGF signalling^40^, we therefore examined whether these pathways regulate OCIAD2 expression.

**Figure 1.**
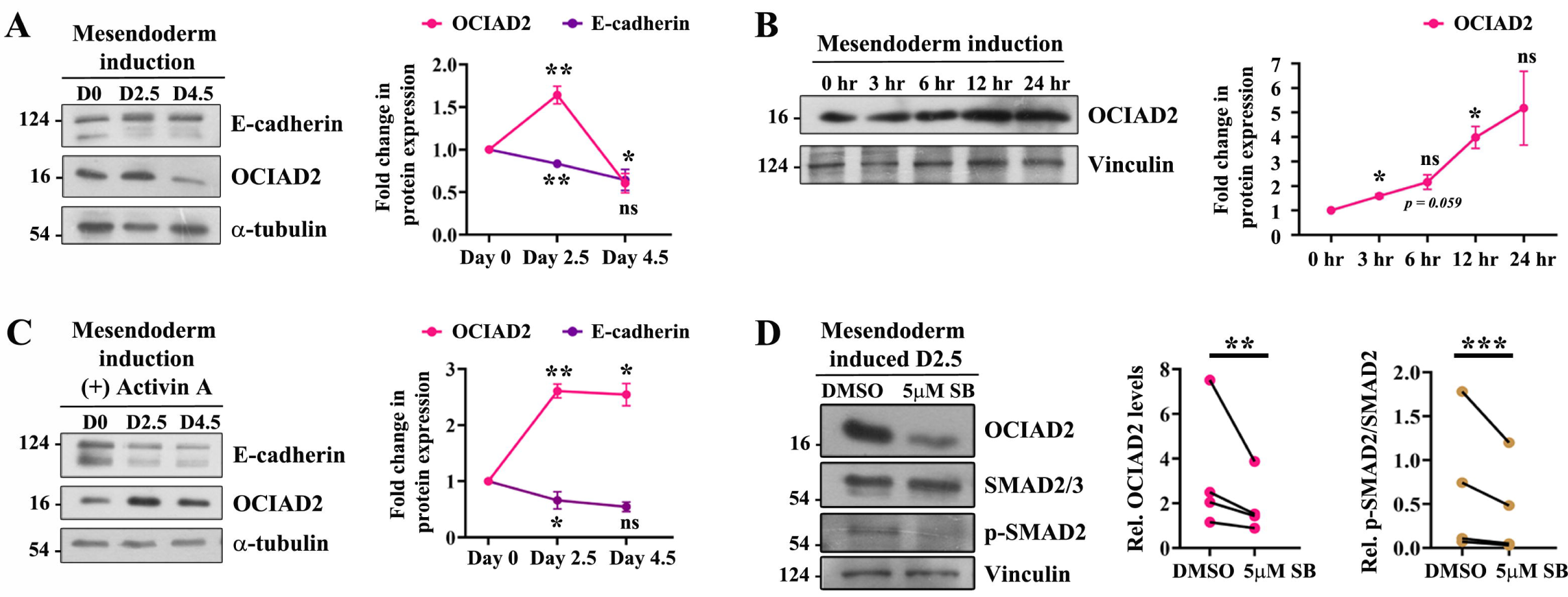
OCIAD2 is induced by Activin A signalling during mesendoderm specification. **(A)** OCIAD2 protein expression kinetics during mesendoderm differentiation. E-cadherin downregulation marks mesendoderm specification. Protein levels were normalized to α-tubulin. **(B)** Immunoblotting analysis for OCIAD2 protein expression upon mesendoderm induction. Graph show quantification of fold change in protein levels normalized to vinculin. **(C-D)** Immunoblotting analysis of OCIAD2 levels upon **(C)** prolonged Activin A treatment during mesendoderm induction from day 0 to day 4.5 or **(D)** inhibition of Activin A signalling using 5μM SB431542 (SB) from day 0 to day 2. Graphs show quantification of fold change in protein levels (for **C**) or relative protein levels (for **D**), normalized to α-tubulin or vinculin, respectively. Results shown are a representative of three independent experiments. Statistical significance was calculated using one sample two-tailed t-test with respect to 0 hr (hour) sample (**A-C**) and ratio paired two-tailed t-test (**D**). Error bars denote standard error of mean, \**p*<0.05, \*\**p*<0.01, \*\*\**p*<0.001, ns indicates non-significant.

### OCIAD2 expression depends on TGF**β**/Activin A signalling

High throughput studies in hESCs reported that TGFβ stimulation leads to SMAD3 binding at the *OCIAD2* locus^41, 42^, suggesting that TGFβ may regulate OCIAD2 levels. Protein pathway prediction studies in cancer cells^30^ and expression analysis in immortalized keratinocytes^43^ have also suggested that TGFβ stimulation can upregulate OCIAD2. To test whether *OCIAD2* expression in hPSC is dependent on TGFβ signalling, we treated hESC cultures with SB431542 (SB), a TGFβ type I receptor blocker. There was a 50% reduction in *OCIAD2* transcript levels, confirming that OCIAD2 is a downstream target of TGFβ signalling in hESCs. Transcript levels of *NANOG* and *CDH1*, direct targets of the TGFβ pathway, were also reduced (Figure S2C).

TGFβ/Activin A and BMP signalling pathways play essential roles in axis patterning and morphogenesis during embryonic development^44^ and in driving mesendoderm differentiation *in vitro*^45, 46^. The dynamic expression profile of OCIAD2, seen upon mesendoderm induction (Figure 1A), positively correlated with the availability of Activin A in the differentiation regime (Figure S2A). As the differentiation protocol involves the activation of two TGFβ superfamily signalling pathways-TGFβ/Activin A and BMP, we investigated their roles in regulating OCIAD2 expression. To test whether Activin A is responsible for increased OCIAD2 expression, we continued Activin A supplementation till day 4.5 and found sustained high OCIAD2 expression levels (Figure 1C). Conversely, inhibition of Activin A signalling with SB431542 significantly reduced OCIAD2 expression (Figure 1D; Figure S2A). Blocking BMP4 signalling with Dorsomorphin (DM), a BMP type I receptor inhibitor, had no effect on OCIAD2 levels (Figure S2A; Figure S2D-F). Collectively, our results demonstrate that TGFβ/Activin A signalling, and not BMP4, induces OCIAD2 expression in hESCs during mesendoderm differentiation.

### Elevated OCIAD2 levels promote EMT in hESCs

Our data shows that OCIAD2 is expressed in undifferentiated hESCs and its expression steadily increases as differentiation progresses, in a TGFβ/Activin A-dependent manner. To test the impact of OCIAD2 levels on hESC maintenance and differentiation, we generated OCIAD2 depleted (knockout: KO) and overexpressing (OV) hESC lines. Analysis for pluripotency-associated marker gene expression, as well as spontaneous trilineage differentiation, showed that OCIAD2 depletion or overexpression had no apparent impact on pluripotency or germ layer specification, suggesting that OCIAD2 modulated hESCs maintain their stem cell identity^33, 34^. Further, OCIAD2 modulation did not impair cell proliferation or induce apoptosis in undifferentiated hESCs (Figure S3A-C).

To investigate whether OCIAD2 modulation influences gene expression in PSCs, we performed a bulk mRNA transcriptome (RNA-seq) analysis. While OV had a modest impact on the transcriptome [52 DEGs (differentially expressed genes)], 661 genes (460 downregulated, 201 upregulated) were differentially expressed in KO hESCs (Figure 2A-B). Importantly, OCIAD2 modulation (KO or OV) did not alter the expression of key pluripotency-associated genes, aligning with our previous findings that OCIAD2 is not essential for maintaining hESC pluripotency^33, 34^.

**Figure 2.**
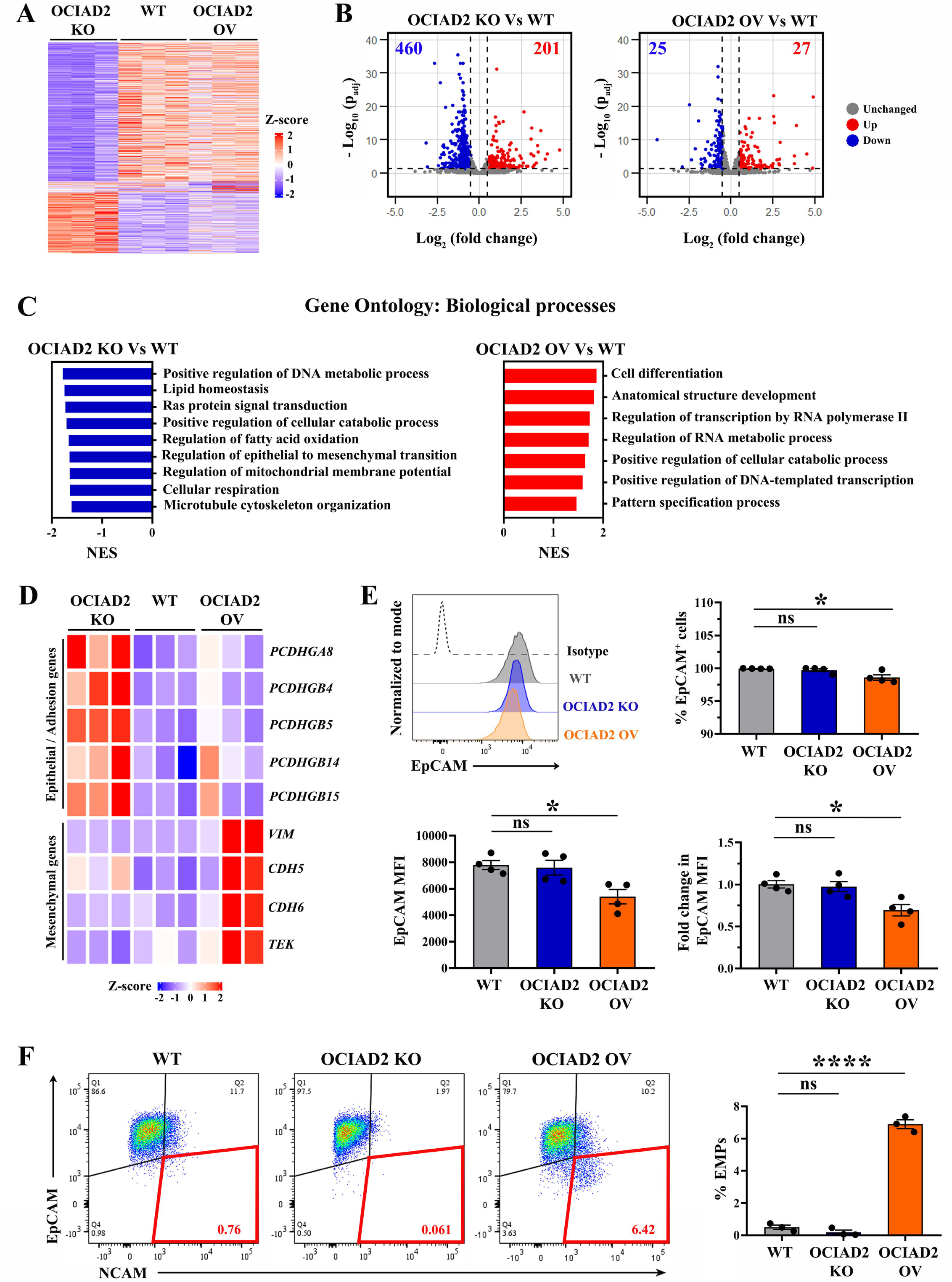
OCIAD2 overexpressing hESCs show hallmarks of EMT. **(A)** Heatmap analysis of WT, OCIAD2 KO and OCIAD2 OV RNA-seq data representing all significant DEGs [log_2_ FC > ± 0.5; p_adj_ value < 0.05]. **(B)** Volcano plots depicting DEGs in OCIAD2 KO or OCIAD2 OV hESCs. Points in red and blue are significantly (-log_10_ p_adj_ > 1.3) upregulated (log_2_ FC > 0.5) and downregulated (log_2_ FC < - 0.5) genes, respectively. **(C)** Bar graphs representing Gene Ontology (GO) analysis of biological processes enriched in day 0 OCIAD2 KO and OCIAD2 OV hESCs, represented by the Normalized Enrichment Score (NES). Positive NES values (red bars) indicate upregulation, whereas negative NES values (blue bars) signify downregulation. **(D)** Heatmap depicting the expression of epithelial/adhesion and mesenchymal genes in day 0 OCIAD2 modulated hESCs. **(E)** Flow cytometry analysis of EpCAM expression in day 0 OCIAD2 modulated hESCs, represented by histogram overlay. Histogram with the dotted line represents the isotype control. Graphs show quantification of % EpCAM expressing cells, EpCAM MFI (median fluorescence intensity) and fold change in EpCAM MFI levels. **(F)** Flow cytometry analysis of EpCAM^-^ NCAM^+^ (EMPs) at day 0, confirming the onset of EMT in OCIAD2 OV hESCs. Gates outlined in red mark EMPs. Results shown are a representative of at least three independent experiments. One-way ANOVA with Dunette’s multiple comparisons test (for total EpCAM^+^ cells, EpCAM MFI and % EMPs) or one sample two-tailed t-test (for fold change in EpCAM MFI) was used for statistical analysis. Error bars denote standard error of mean, \**p*<0.05, \*\*\*\**p*<0.0001, ns indicates non-significant.

Gene ontology (GO) analysis of DEGs in OCIAD2 KO hESCs revealed upregulation of biological processes such as nucleosome assembly, chromatin remodelling, protein-DNA complex organization and calcium dependent cell-cell interactions (Figure S3D), while epithelial to mesenchymal transition (EMT), cytoskeleton organization, Ras signal transduction, mitochondrial membrane potential and several metabolic processes (lipid homeostasis, DNA metabolism and fatty acid oxidation) were downregulated (Figure 2C). Pathway enrichment analysis highlighted that genes associated with focal adhesion, ECM-receptor organization, amino acid (valine, leucine and isoleucine) and propanoate metabolism were perturbed upon loss of OCIAD2 (Figure S3E). GO analysis of DEGs from OV hESCs showed enrichment only in upregulated processes, which were related primarily to cell differentiation, pattern specification, transcription and metabolism (Figure 2C). Moreover, Gene Set Enrichment Analysis (GSEA) of OV DEGs showed positive enrichment of TGFβ and WNT signalling (Figure S3F), pathways that are essential for early lineage specification^47^. Collectively, this suggests that increased OCIAD2 expression may render hESCs more prone to differentiation.

Pathway prediction analysis has previously suggested that TGFβ-induced OCIAD2 expression may promote EMT in tumors^30^. Since KO hESCs showed downregulation of processes such as EMT and cytoskeleton organization, we asked whether OCIAD2 levels regulate EMT in hESCs. Expression analysis of epithelial and mesenchymal gene signatures revealed that OCIAD2 KO hESCs showed increased expression of protocadherin genes, which are associated with cell-cell adhesion, whereas OCIAD2 OV hESCs exhibited upregulation of mesenchymal markers such as *VIM*, *CDH5*, *CDH6*, and *TEK* (Figure 2D). Flow cytometry analysis showed that OV hESCs displayed only a marginal reduction in the percentage of cells expressing the epithelial marker EpCAM (Epithelial Cell Adhesion Molecule), with expression levels reduced by about 25% (Figure 2E). Concomitantly, we observed that NCAM-1 (Neural Cell Adhesion Molecule-1), an early mesenchymal marker was upregulated in cells with reduced EpCAM levels (Figure 2F). EpCAM^-^ NCAM^+^ cells, referred to as early mesendoderm progenitors (EMPs) typically arise after about 3.5 days of mesoderm induction of hESCs^40^. The presence of EMPs in uninduced OV cultures suggests that OCIAD2 overexpression may predispose hESCs towards mesendoderm differentiation.

### Loss of OCIAD2 delays early mesendoderm specification

Mesendoderm differentiation involves an epithelial to mesenchymal transition (EMT) and is characterized by the downregulation of epithelial markers such as E-cadherin, the upregulation of mesenchymal markers such as N-cadherin, Snail and Twist, and extensive remodelling of the ECM and cytoskeleton^47, 48^. Since OCIAD2 levels affect the expression of EMT markers, we posited that OCIAD2 may regulate developmental EMT during early lineage specification.

Immunostaining analysis at day 3.5 of differentiation, when EMPs first appear, showed that OCIAD2 KO had reduced vimentin or N-cadherin expression, while OV cells expressed higher levels of these markers (Figure 3A-B). This correlated with elevated transcript levels of the mesenchymal markers *SNAI2* and *VIM* at day 3.5 (Figure 3C), indicating that increased OCIAD2 expression favors EMT.

**Figure 3.**
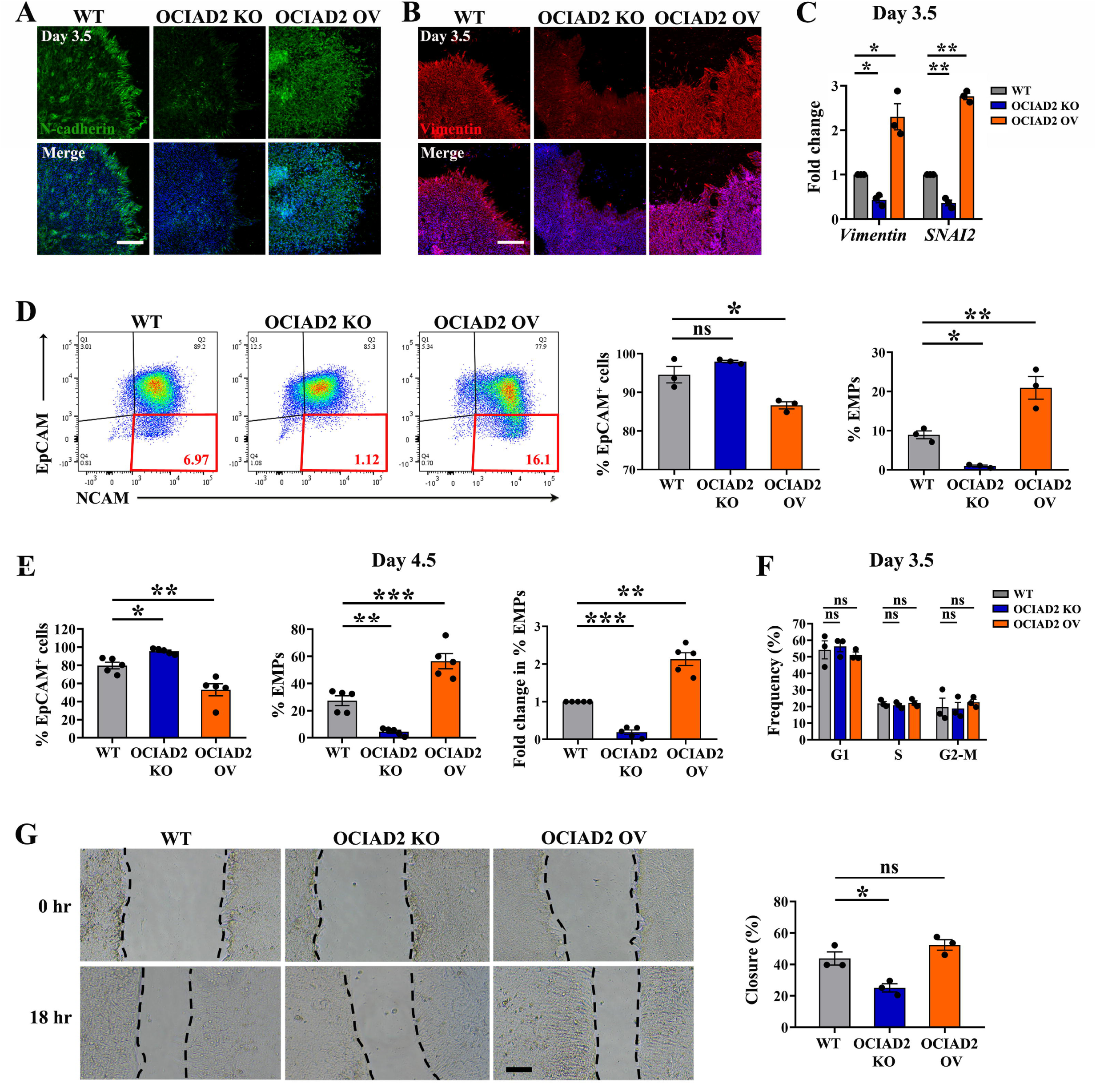
OCIAD2 regulates EMT during mesendoderm specification of hESCs. **(A-B)** Immunostaining of mesendoderm-induced day 3.5 cultures with mesenchymal markers **(A)** N-cadherin and **(B)**Vimentin. Nucleus is marked using DAPI. Scale bar: 100μm. **(B)** RT-qPCR analysis of EMT markers *SNAI2* and *VIM* in mesendoderm-induced cells at day 3.5. Transcript levels normalized to RPLP0. **(D-E)** Flow cytometry analysis of EpCAM^-^ NCAM^+^ cells (EMPs) at **(D)** day 3.5 and **(E)** day 4.5. Gates outlined in red mark EMPs. Graphs show quantification of % EpCAM^+^ cells, % EMPs and fold change in % EMPs. **(F)** Proliferation and cell cycle analysis of mesendoderm-induced day 3.5 cultures using Ki-67/Hoechst 33342 staining. Boxed regions denote the different cell cycle stages identified using Hoechst 33342 staining. Bar graph represents frequency of these cell cycle stages in mesendoderm-induced day 3.5 OCIAD2 modulated cultures. **(G)** Representative phase contrast microscopy images of scratch wound closure of mesendoderm-induced cultures at 0 hr and 18 hrs post-scratching. Dotted black lines represent the scratched edges. Scale bar: 50μm. Graph represents quantification of percentage of scratch wound closure after 18 hrs. A minimum of three scratches were measured from each condition, from three independent experiments. Results shown are a representative of at least three independent experiments. One-way ANOVA with Dunette’s multiple comparisons test (**D-G**) or one sample two-tailed t-test [for **(A) (C)** and fold change comparisons in **(E)**] was used for statistical comparison. Error bars denote standard error of mean, \**p*<0.05, \*\**p*<0.01, \*\*\**p*<0.001, ns indicates non-significant.

Flow cytometry analysis of induced cultures at day 3.5 showed that compared to WT (8.9%), KO cells generated significantly fewer EMPs (1.3%), whereas OV had an approximate 2-fold increase in EMP production (20.9%). Consistent with these findings, there was a marked reduction in EpCAMC OV cells (86.6%) compared with WT cells (94.5%). The majority of KO cells (98.5%) remained EpCAM^+^, indicating that they retain their epithelial identity (Figure 3D). At day 4.5, KO cells (95.5%) continued to express EpCAM and generated few EMPs (4.5%), indicative of incomplete EMT. In contrast, there was robust differentiation of OV cells to EMPs (55.8%) (Figure 3E).

Flow cytometry analysis of Ki-67 and Hoechst 33342 stained cultures at 3.5 days showed that there was no significant difference in the distribution of cells across the G1, S, or G2-M phases in KO or OV cells, confirming that OCIAD2 has no gross effect on cell proliferation (Figure 3F). A scratch wound healing assay showed that while WT cells and OV cells had a comparable extent of wound closure in the assay (WT: 45%; OV: 52%), KO showed only 25% closure (Figure 3G), suggesting that optimal levels of OCIAD2 are essential for migration of early mesendoderm derivatives.

While the employed mesendoderm differentiation regimen relies on minimal extrinsic signals (Activin A, BMP4, VEGF and FGF2) to induce the PS, additional WNT activation in the same context robustly generates PS-like populations within 24 hrs^39^. To test whether KO cells can differentiate in strong inducing conditions, we supplemented the differentiation media with CHIR99021 (WNT agonist). This resulted in complete downregulation of EpCAM, with no significant differences in the percentages of EMPs between WT and KO hESCs at day 4 (Figure S4A). Similarly, KO hESCs could efficiently generate EMPs when subjected to a commercial mesoderm induction media (STEMdiff^TM^) (Figure S4B). These findings suggest that sustained induction with CHIR99021 may potentially mask the effects of OCIAD2 on mesendoderm specification, and more importantly, OCIAD2 may function in coordination with signalling pathways such as WNT to orchestrate lineage commitment.

To test whether OCIAD2 affects ectoderm specification, we performed neuroectoderm differentiation. KO cells showed reduced Pax6^+^ neural cells, whereas OCIAD2 overexpression promoted the formation of Pax6^+^ clusters (Figure S4C), indicating that OCIAD2 also enhances neuroectoderm differentiation. Collectively, our results demonstrate that while OCIAD2 loss does not entirely abolish differentiation capabilities, OCIAD2 is required for efficient differentiation of hESCs.

### OCIAD2 depletion dysregulates TGF**β**, BMP, PI3K-AKT and MAPK/ERK pathways to influence mesendoderm fate

Since OCIAD2 expression is upregulated at early stages of mesendoderm induction and peaks by day 2, we investigated whether it has a role in initiating EMT. Transcriptome analysis at day 2 identified 1068 and 395 DEGs in KO and OV, respectively (Figure S5A). Key biological processes downregulated in KO cells included developmental processes such as anterior-posterior axis specification, EMT, and symmetry specification. Additionally, processes related to transcription, transcriptional regulation, RNA processing and serine phosphorylation were suppressed, highlighting the broader impact of OCIAD2 depletion on gene expression during mesendoderm differentiation. Further, downregulation of mitochondrial transport, fatty acid catabolism and nucleic acid metabolism, suggested effects on cellular metabolism, as was also observed in the day 0 RNA-seq analysis. Conversely, all of the aforementioned pathways were upregulated in mesendoderm-induced OV cells, with enrichment of GO terms such as gastrulation, endoderm differentiation, stem cell differentiation and EMT (Figure 4A). Altogether, this indicates that the impact of OCIAD2 levels on differentiation begins as early as day 2, suggesting a pivotal role in driving mesendoderm specification.

**Figure 4.**
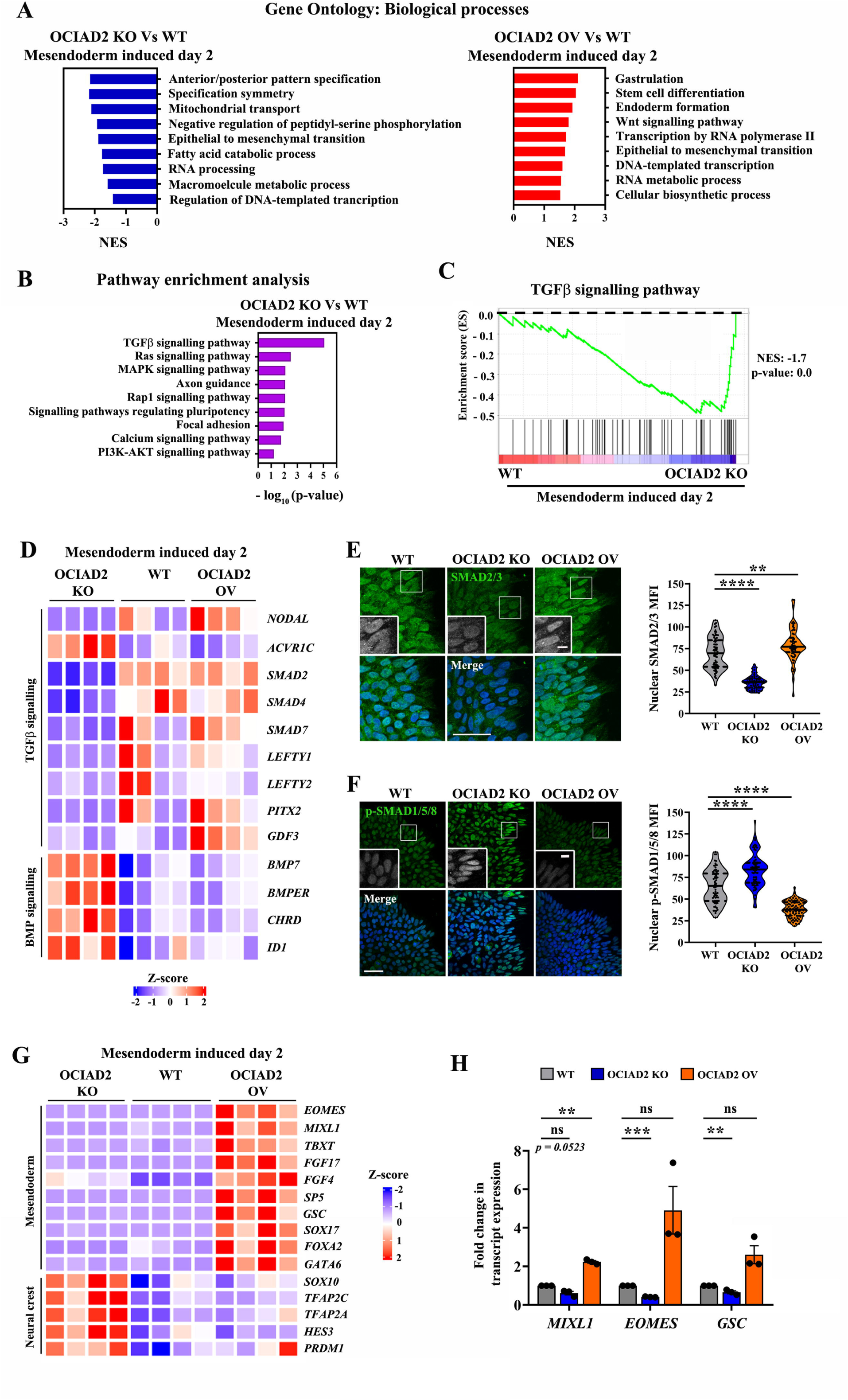
OCIAD2 regulates early mesendoderm specification via controlling TGFβ/BMP and WNT signalling pathways. **(A)** Bar graphs representing Gene Ontology (GO) analysis of biological processes enriched in mesendoderm-induced day 2 OCIAD2 KO and OCIAD2 OV precursors, represented by the Normalized Enrichment Score (NES). Positive NES values (red bars) indicate upregulation, whereas negative NES values (blue bars) signify downregulation. **(B)** Bar graph showing top significantly enriched pathways identified using KEGG pathway enrichment analysis in OCIAD2 KO and early mesendoderm precursors at day 2. **(C)** Gene set enrichment analysis (GSEA) plot for TGFβ pathway genes in mesendoderm-induced day 2 WT and OCIAD2 KO cells. NES value indicates pathway enrichment. p-value = 0.0 indicates statistically significant. **(D)** Heatmap depicting the expression of TGFβ and BMP pathway genes in mesendoderm-induced OCIAD2 modulated lines at day 2. **(E-F)** Immunostaining of **(E)** total SMAD2/3 in mesendoderm-induced cultures 6 hrs post-induction and **(F)** phosphorylated SMAD1/5/8 (p-SMAD1/5/8) in mesendoderm-induced cultures 3 hrs post-induction. Boxed regions are magnified and shown as insets. Violin plots show the quantification of **(E)** nuclear SMAD2/3 and **(F)** nuclear p-SMAD1/5/8 accumulation. DAPI staining was used to identify the nuclear area. Each data point represents a single nucleus, with at least 20 cells measured from three independent experiments. Scale bar: 50μm; 10μm (insets). **(G)** Heatmap representing the expression of mesendoderm and neural crest progenitor (NCp) genes at day 2 of mesendoderm induction. **(H)** RT-qPCR analysis of early mesendoderm markers in OCIAD2 modulated lines after 6 hrs of induction. Transcript levels normalized to RPLP0. Results shown are a representative of at least three independent experiments. One-way ANOVA with Dunette’s multiple comparisons test (for **E-F**) or one sample two-tailed t-test (for **H**) was used for statistical comparison. Error bars denote standard error of mean, \**p*<0.05, \*\**p*<0.01, \*\*\*\**p*<0.0001, ns indicates non-significant.

Pathway enrichment analysis identified TGFβ signalling as the most enriched pathway in differentiating OCIAD2 KO cells (Figure 4B). Gene set enrichment analysis (GSEA) further confirmed negative enrichment of TGFβ pathway genes in KO mesendoderm precursors, indicating that OCIAD2 depletion suppresses TGFβ pathway activity in mesendoderm-induced hESCs (Figure 4C). Notably, several TGFβ cascade genes such as *NODAL*, *SMAD2*, *SMAD4*, *SMAD7*, *LEFTY1*, *LEFTY2* and *PTIX2* were downregulated. In contrast, *ACVR1C*, a type I TGFβ receptor, was markedly upregulated (Figure 4D). Immunostaining for total SMAD2/3 after 6 hrs of mesendoderm induction showed diminished nuclear accumulation of SMAD2/3 in KO, while OV cells exhibited enhanced nuclear SMAD2/3 levels (Figure 4E). These findings indicate that OCIAD2 is required for effective activation of the TGFβ signalling pathway, and that its loss results in transcriptional repression of key pathway effectors.

While the expression of core BMP signalling components was not significantly altered, we detected elevated transcripts for BMP signalling targets such as *BMP7*, *BMPER* and *CHRD* in mesendoderm-induced KO cells (Figure 4D). This suggests that autocrine BMP signalling may be enhanced in the absence of OCIAD2. Immunostaining for phosphorylated SMAD1/5/8 (p-SMAD1/5/8), a hallmark of active BMP signalling, revealed increased nuclear localization in KO, indicating elevated BMP signalling activity. Conversely, OV had reduced nuclear p-SMAD1/5/8 levels, indicating attenuation of the pathway (Figure 4E). Taken together, our data demonstrate that OCIAD2 exerts opposing effects on the TGFβ and BMP signalling cascades.

In addition to TGFβ signalling, genes associated with PI3K/AKT, MAPK/ERK and Ras signalling were also enriched in mesendoderm-induced KO cells (Figure 4B). While our day 0 RNA-seq analysis did not report these signalling cascades, several processes dependent on these pathways, such as cell-cell adhesion, ECM-receptor interaction, focal adhesion were enriched in KO hESCs (Figure 1C; Figure S3E). Moreover, we observed downregulation of core TGFβ signalling components such as *TGFA*, *TGFB2*, *SMAD2*, *SMAD4*, *TGIF*, *NEDD4L* in KO hESCs (Figure S5C). This suggests that OCIAD2 depletion attenuates TGFβ signalling in undifferentiated hESCs as well. TGFβ/SMAD cooperates with PI3K-AKT pathway to maintain self-renewal, whereas MAPK/ERK and WNT promote differentiation in hESCs^49^. Since our experimental design involves modulating OCIAD2 levels in undifferentiated hESCs, we hypothesized that the observed effects on signalling and lineage outcomes are initiated prior to mesendoderm induction. Therefore, we chose to assess the activation of TGFβ and PI3K-AKT signalling at the undifferentiated stage (day 0), as they are known to coordinate SMAD2/3 activity and balance self-renewal versus mesendoderm specification. Expectedly, KO hESCs exhibit reduced total SMAD2 and phosphorylated SMAD2 (p-SMAD2) levels, whereas OV showed increased p-SMAD2 levels (Figure S5D). Further, KO hESCs showed reduced total AKT levels, suggesting possible transcriptional downregulation; however, this was not accompanied by a change in total phosphorylated AKT (p-AKT) levels, indicating compensatory activation. In contrast, OV hESCs maintained total AKT levels but showed reduced p-AKT (Figure S5E). This reduction in p-AKT, alongside elevated p-SMAD2/3, may impair self-renewal and initiate differentiation. Collectively, these findings suggest that OCIAD2 modulates TGFβ and PI3K-AKT signalling thresholds to regulate early cell fate decisions in hESCs.

### OCIAD2 promotes the activation of early mesendoderm genes

The coordinated activation of BMP, WNT and TGFβ/Activin A/Nodal signalling facilitates the downregulation of pluripotency genes and enhances the transcription of early mesendoderm targets, thereby establishing lineage commitment^50, 51^. We hypothesized that aberrant activation of these signalling pathways in KO hESCs could hinder the acquisition of the mesendodermal fate, ultimately restricting differentiation. To investigate this, we analysed the expression of early lineage-associated genes upon mesendoderm induction at day 2. Mesendoderm-induced OV cells showed increased expression of mesodermal markers, such as *MIXL1*, *TBXT*, *SP5*, *FGF4*, *FGF17* as well as endodermal genes, including *EOMES*, *GSC, SOX17* and *FOXA2*, indicating an accelerated response to induction. On the other hand, expression levels of mesendoderm genes were statistically comparable between WT and KO mesendoderm precursors (Figure 4G). We reasoned that this similarity might be due to the timing of the transcriptomic analysis, which was conducted on day 2, when signalling responses may already be attenuated. To assess the immediate effects of OCIAD2 modulation on acquisition of mesendoderm fate, we performed RT-qPCR analysis of mesendoderm genes six hrs post-induction. We found reduced *EOMES* and *GSC* transcript levels and a downward trend in *MIXL1* expression in KO, whereas OV cells showed an increasing trend for *GSC* and *EOMES* expression, along with a significant upregulation of *MIXL1* (Figure 4H). These results indicate that OCIAD2 facilitates the induction of mesendoderm genes.

Interestingly, KO cells showed increased expression of a subset of neural crest progenitor (NCp) genes, specifically *SOX10*, *TFAP2A*, *TFAP2C, HES3* and *PRDM1* (Figure 4G). During early phases of mesendoderm induction, WNT-mediated stabilization of β-catenin promotes the transcription of NCp genes. The synergistic interactions between SMAD2/3 and β-catenin inhibit NCp gene expression and initiate transcription of PS genes, thereby stabilizing mesendoderm commitment^52^. The upregulation of NCp markers in mesendoderm-induced KO cells suggests a block in lineage progression, likely due to insufficient SMAD2/3 activity. Thus, it is plausible that KO cells fail to efficiently commit to the mesendoderm lineage as a result of impaired TGFβ signalling. Taken together, these findings suggest that OCIAD2 is potentially required for TGFβ-mediated activation of the mesendodermal gene program.

### OCIAD2 controls mitochondrial morphology in hESCs

OCIAD2 is a mitochondrial protein previously reported to interact with the electron transport chain (ETC) proteins^22, 28^. Additionally, loss of OCIAD2 disrupts gross mitochondrial structure and induces apoptosis in lung adenocarcinoma cell lines^29^. Hence, we investigated whether OCIAD2 may regulate mesendoderm specification through mitochondrial function. In undifferentiated hESCs, mitochondria are typically short and spherical, displaying perinuclear clustering^53, 54^. Visualization of the mitochondrial network using Mitotracker deep red, showed elongated mitochondria with increased median branch length in KO hESCs. However, the mitochondrial network was extensively hyperfused resulting in a reduced mitochondrial footprint (cellular area covered by mitochondrial structures) and a possible shift towards increased fusion (Figure 5A; Video S1). Live imaging showed significantly lower mitochondrial dynamics (measured as variance of the mitochondrial network parameters-number of branches, junctions, and mitochondrial footprint over time) in KO hESCs compared to WT cells (Figure S6A; Video S1). This suggests that the loss of OCIAD2 perturbs mitochondrial network architecture in hESCs.

**Figure 5.**
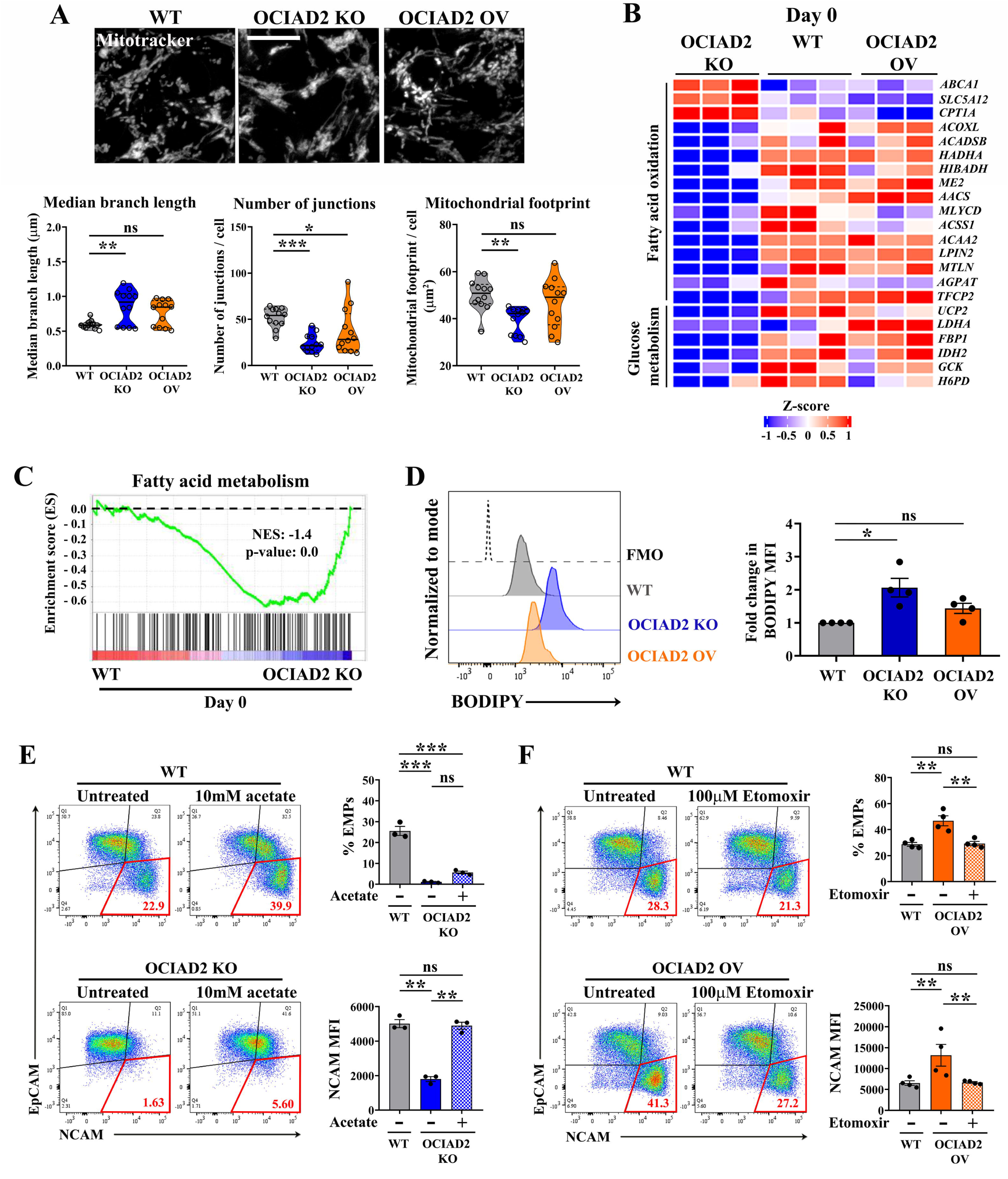
OCIAD2 influences fatty acid oxidation to regulate EMT. **(A)** 3D projected images of MitoTracker deep red labelled mitochondria in OCIAD2 modulated hESCs. Scale bar: 10μm. Violin plots showing quantification of mitochondrial branch length, number and footprint. At least 50 cells from three replicates were analysed. **(B)** Heatmap showing DEGs enriched in glucose metabolism and fatty acid oxidation in from day 0 RNA-seq. **(C)** GSEA plot for fatty acid metabolism genes in day 0 OCIAD2 KO hESCs. NES value indicates pathway enrichment. p-value = 0.0 indicates statistically significant. **(D)** Histogram overlay of BODIPY staining representing increased accumulation of neutral lipids in OCIAD2 KO hESCs. Histogram with the dotted line represents the fluorescence minus one (FMO) control. Graph shows quantification of fold change in BODIPY MFI. **(E-F)** Flow cytometry analysis of EpCAM^-^ NCAM^+^ cells (EMPs) at day 3.5 in **(E)** WT and OCIAD2 KO hESCs treated with 10mM sodium acetate and **(F)** WT and OCIAD2 OV treated with 100μM etomoxir. Gates outlined in red mark EMPs. Graphs show quantification of % EMPs and NCAM MFI. Results shown are a representative of at least three independent experiments. Statistical significance was calculated using Kruskal Wallis test (for **A**), or one sample two-tailed t-test (for **D**) or two-way ANOVA with Tukey’s multiple comparisons test (for **E-F**). Error bars denote standard error of mean, \**p*<0.05, \*\**p*<0.01, \*\*\*\**p*<0.0001, ns indicates non-significant.

### OCIAD2 depletion perturbs lipid metabolism in hESCs

hESCs primarily rely on glycolysis but transition to oxidative phosphorylation (OXPHOS) during mesendoderm differentiation^9, 53, 55, 56^. OCIAD2 interacts with Complex I, III and IV and controls mitochondrial bioenergetics in HEK293 cells^28^. Despite alterations in mitochondrial morphology, subjecting hESCs to a mito stress assay to assess mitochondrial respiration, showed no significant differences in the oxygen consumption rates upon OCIAD2 modulation (Figure S6B).

RNA-seq analyses at day 0 and day 2 consistently showed enrichment of pathways related to lipid homeostasis, fatty acid oxidation (FAO), propanoate metabolism, and the catabolism of branched-chain amino acids (BCAAs), including valine, leucine, isoleucine, and β-alanine in KO hESCs (Figure 1C; Figure S3C; Figure 4A; Figure S5B). Analysis of the DEGs in KO hESCs showed upregulation of plasma membrane fatty acid transporters *ABCA1* and *SLC5A12*, as well as *CPT1A*, the outer mitochondrial membrane transporter responsible for importing fatty acids into mitochondria. However, several downstream FAO genes were downregulated, suggesting impaired lipid catabolism upon the loss of OCIAD2 (Figure 5B). GSEA further indicated a negative enrichment of fatty acid metabolism upon OCIAD2 depletion (Figure 5C). Additionally, we observed downregulation of genes involved in glucose metabolism, including *UCP2*, *LDHA*, *GCK*, *IDH2*, *GALE* and *H6PD* (Figure 5B). Flow cytometry-based uptake analysis of 2-NBDG, a fluorescent glucose analog, showed that glucose uptake upon OCIAD2 modulation was comparable to WT (Figure S6C). Further, BODIPY staining, which labels neutral lipids, was elevated in KO hESCs (Figure 5D), indicating increased fatty acid accumulation upon OCIAD2 depletion. Thus, OCIAD2 possibly affects glucose and fatty acid metabolism in hESCs, without impacting mitochondrial bioenergetics.

### OCIAD2 controls lipid oxidation to support mesendoderm specification

High-throughput mass spectrometry analysis of the mitochondrial proteome identified that OCIAD2 interacts with Acyl-CoA thioesterase 9 (ACOT9) and Hydroxyacyl-CoA dehydrogenase (HADHA)^57^, enzymes involved in the terminal steps of β-oxidation of long-chain fatty acids. Since OCIAD2 depletion resulted in selective downregulation of FAO genes, we hypothesized that OCIAD2 primarily influences fatty acid oxidation, rather than synthesis, in hESCs. FAO is a catabolic process that breaks down long-chain fatty acids to produce acetyl-CoA, which enters the tricarboxylic acid (TCA) cycle for ATP production^58^.

Hence, it is possible that loss of OCIAD2 may disrupt FAO, leading to reduced acetyl-CoA levels, ultimately impairing mesendoderm specification. To test this, we supplemented sodium acetate, a precursor of acetyl-CoA, during mesendoderm induction. Treatment of WT cells with 10 mM sodium acetate resulted in significant reduction in EpCAM^+^ cells, seen by flow cytometry analysis. This was accompanied by an upregulation of NCAM levels and increased EMPs by day 3.5 (Figure S6D), demonstrating that increased acetate levels enhance mesendoderm specification. Notably, KO hESCs responded to acetate supplementation, showing increased NCAM expression to the extent seen in WT EMP differentiation and improved EMP generation (Figure 5E). However, this did not sufficiently downregulate EpCAM (Figure S6D), therefore only partially restoring mesendoderm differentiation potential in KO.

Inhibition of FAO in WT hESCs using 100μM etomoxir, a CPT1A inhibitor that blocks the import of long chain fatty acids into mitochondria, led to a minor reduction in NCAM expression and EMPs. In contrast, etomoxir treatment of OV hESCs resulted in an increase in EpCAM expressing cells (Figure S6E). This was also accompanied by a marked reduction in NCAM levels and mesendoderm differentiation, thereby restoring EMP frequencies to WT hESCs (Figure 5F). This indicates that OCIAD2 promotes differentiation via mitochondrial FAO. Therefore, increased OCIAD2 expression may render hESCs more dependent on FAO to fuel mesendoderm differentiation. Collectively, our findings point toward a potential role for OCIAD2 in modulating metabolic dependencies that influence lineage differentiation outcomes.

## Discussion

Our study identifies OCIAD2 as a critical regulator of mesendoderm differentiation in human pluripotent stem cells (hPSCs), acting at the intersection of mitochondrial metabolism and signalling. While previous reports have linked OCIAD2 to cancer progression and mitochondrial function^26, 28, 59^, its physiological role in early human development remained unexplored. Here, we demonstrate that OCIAD2 is transiently upregulated during mesendoderm induction in a TGFβ/Activin A-dependent manner, and its depletion impairs mesendoderm lineage specification by disrupting epithelial-to-mesenchymal transition (EMT).

OCIAD2 is a vertebrate gene expressed in early embryogenesis^32, 60^, although the significance of this expression is unknown. We show that upregulation of OCIAD2 is critical for early stages of mesendoderm differentiation, and its depletion impairs lineage specification. Although OCIAD2 depletion or overexpression does not affect pluripotency or trigger aberrant differentiation in hESCs, it alters transcriptional programs toward epithelial (KO) or mesenchymal (OV) states, indicating that OCIAD2 is a gatekeeper of differentiation.

OCIAD2 depletion compromised the induction of mesendoderm genes, resulting in defective specification. Aberrant OCIAD2 expression, along with reduced association with mesenchymal fate, has been reported during zebrafish tail regeneration^61^. OCIAD2 has been identified as a prognostic marker in invasive lung adenocarcinomas and ovarian mucinous tumors^59, 62, 63^. Collectively, these reports indicate that OCIAD2 may likewise regulate EMT in regenerative and metastatic contexts. Although our analysis shows that OCIAD2 modulation dysregulates epithelial/mesenchymal characteristics in hESCs, abnormal phenotypes manifest only after providing differentiation cues. This suggests that OCIAD2 depends on inductive signals to robustly drive developmental EMT. However, whether this is true for other forms of EMT, remains to be tested.

Though previous reports show that TGFβ signalling can upregulate OCIAD2^31, 43^, the underlying mechanisms and its regulation in human pluripotent stem cells was not known. Our findings report that OCIAD2 is transiently upregulated during mesendoderm induction via TGFβ/Activin A signalling, and that its loss transcriptionally attenuates TGFβ pathway. Thus, OCIAD2 may facilitate a feedforward loop in the regulation of TGFβ signalling in hESCs. Interestingly, *ACVR1C*, encoding the activin receptor, is upregulated in KO, suggesting a compensatory response. TGFβ signalling is known to activate non-canonical branches, including the PI3K-AKT, MAPK, and Ras pathways, via its receptors^64^, which may explain the observed disruption of these pathways upon OCIAD2 depletion. Although TGFβ signalling promotes pluripotency by directly regulating Nanog^41^, KO hESCs maintain pluripotency despite reduced SMAD2/3 activation but fail to initiate mesendoderm differentiation. During gastrulation, transient TGFβ/Activin A activation sustains pluripotency but is insufficient to drive mesendoderm specification^65^. This transient activation maintains a basal signalling threshold, such that responses to further ligand stimulation are minimal. In contrast to prolonged ligand exposure, repeated pulses of TGFβ/Activin A enhance primitive streak differentiation in a 2D gastruloid model^66^. Additionally, SMAD proteins exhibit differential affinities for their target genes that may enable TGFβ/Activin A signalling to distinguish between transcriptional programs for pluripotency and mesendoderm specification^67^. Thus, reduced SMAD2/3 activity seen in KO hESCs, appears sufficient to maintain pluripotency but is inadequate for differentiation. This suggests that OCIAD2 may be required not merely for sustaining baseline SMAD signalling, but also for modulating the signalling amplitude or dynamics necessary to trigger mesendoderm specification.

Although OCIAD2 expression relies solely on TGFβ/Activin A/Nodal signalling, its loss leads to enhanced activation of BMP4 signalling during early mesendoderm differentiation. Given that SMAD2 downregulation can relieve repression of BMP4 signalling^68^, the reduced SMAD2/3 levels in KO cells likely favor SMAD1/5/8-mediated BMP activation. Therefore, OCIAD2 may orchestrate the cross-regulation between TGFβ and BMP4 signalling cascades. During gastrulation, opposing Nodal and BMP gradients establish the dorsoventral axis, while graded Nodal signalling determines mesoderm versus endoderm fate^44, 69, 70^. Given its role in activating TGFβ superfamily pathways, it will be interesting to investigate the role of OCIAD2 in axis patterning and morphogenesis.

The PI3K/AKT pathway plays a critical role in determining whether TGFβ/Activin A signalling supports self-renewal or initiates differentiation. AKT regulates phosphorylated SMAD2/3 thresholds, where lower levels are permissive for self-renewal whereas loss of AKT signalling enhances SMAD2/3 activity and promotes differentiation^49^. Although OCIAD2 affects AKT activation in carcinomas^26, 27^, we find that OCIAD2 loss reduces total AKT levels without affecting total phosphorylated AKT levels, while its overexpression downregulates AKT phosphorylation in hESCs. This suggests that OCIAD2 exerts a distinct and complex regulatory effect on TGFβ and PI3K/AKT pathways in hESCs, extending beyond mere pathway activation.

WNT/β-catenin signalling controls early lineage specification by directly regulating both primitive streak (PS) and neural crest progenitor (NCp) genes^52, 71^. The synergistic interaction between β-catenin and SMAD2/3 during early stages of mesendoderm induction represses the transcription of NCp genes while promoting PS gene activation, thereby sustaining mesendoderm differentiation^52, 72^. The upregulation of NCp genes in differentiating KO cells, alongside downregulation of PS genes, suggests that OCIAD2 depletion may disrupt the regulatory interplay between β-catenin and SMAD2/3, leading to impaired activation of the mesendoderm program. Taken together, our findings demonstrate that OCIAD2 loss disrupts the collaborative regulation of TGFβ, AKT, BMP and WNT pathways, impairing hESC differentiation.

Emerging studies highlight the non-canonical role of metabolism in influencing stem cell fate by acting upstream of signalling pathways, particularly through post-translational modifications of WNT, TGFβ and ERK signalling components^12–15, 73^. OCIAD2 depletion in hESCs leads to the accumulation of neutral lipids and downregulation of fatty acid oxidation (FAO) genes, indicating a metabolic imbalance. Although FAO is dispensable for pluripotency maintenance^74^, it enhances endoderm differentiation in hESCs by promoting SMAD3 acetylation^73^. hESCs exhibit low fatty acid and cholesterol uptake and primarily generate acetyl-CoA from glycolysis^75^. Indeed, pharmacological enhancement of acetyl-CoA, an end product of FAO, using sodium acetate promoted EMT in WT hESCs. However, this only partially rescued mesendoderm differentiation in KO hESCs. In contrast, inhibition of FAO using etomoxir did not affect EMP generation in WT hESCs but significantly abrogated mesendoderm specification in OV hESCs. Acetate supplementation broadly replenishes the cytosolic acetyl-CoA pool, whereas etomoxir selectively blocks mitochondrial acetyl-CoA production. Given that OCIAD2 interacts with enzymes involved in the terminal steps of FAO^57^, our data position it at the intersection of metabolism and signalling networks that promote EMT and lineage commitment. While the mechanisms by which OCIAD2 shapes metabolic flux merits further investigation, its dual function may explain its frequent dysregulation in cancers, where EMT and metabolic reprogramming are hallmarks of tumor progression.

Loss of OCIAD1 in hESCs results in increased mesendoderm differentiation upon induction^24^. In striking contrast, our results show that loss of OCIAD2 retards mesendoderm differentiation, indicating that OCIAD proteins exert opposing effects on differentiation. A recent proteome and lipidome analysis found that proteins related to FAO are upregulated upon OCIAD1 depletion^76^. Our RNA-seq analysis at day 0 reports downregulation of FAO genes in OCIAD2-deficient hESCs. Intriguingly, the two vertebrate OCIAD family members potentially have divergent effects on lipid metabolism. While differences in interaction partners, or tissue-specific expression patterns may partly explain non-redundant functions, our analyses point toward a more refined regulatory framework governing these proteins.

Strategies to enhance TGFβ signalling for efficient hESC differentiation suggest using ATP and N-acetylcysteine (NAC)^11^, or small-molecule SMAD activators^77^. These approaches reduce reliance on Activin A, cutting costs while maintaining robust differentiation yields. Since these molecules do not directly control TGFβ signalling, off-target effects are expected. Mechanistic insights into the TGFβ-OCIAD2 axis further may uncover novel small molecule modulators that more precisely control TGFβ signalling, offering specificity for directing mesendoderm differentiation.

In summary, we identify the functional relevance of OCIAD2 in early embryonic development, using hESC differentiation as a model. We demonstrate that OCIAD2 is both a downstream target and a positive regulator of TGFβ signalling in hESCs. We find that OCIAD2 controls developmental EMT to drive mesendoderm specification in hESCs, without compromising stem cell identity. Additionally, we identify OCIAD2 as a potential molecular link between TGFβ signalling and mitochondrial FAO, a connection that merits further mechanistic investigation. We propose that OCIAD2 may act as a functional bridge between metabolic regulation and developmental signalling pathways. A deeper understanding of this regulatory network may provide means for improving the efficiency and fidelity of *in vitro* mesendoderm differentiation.

## Experimental procedures

### Human embryonic stem cell culture

The hESC lines-BJNhem20^36^ (hPSCreg identifier: JNCSRe002-A), OCIAD2 KO^33^ (hPSCreg: identifier JNCSRe002-A-1) and OCIAD2 OV^34^ (hPSCreg: identifier JNCSRe002-A-3) were cultured as described previously^36, 78^. Briefly, cells were mechanical passaged every 4 days onto mitomycin treated mouse embryonic fibroblasts (MEFs) in hESC media containing Knockout Dulbecco’s modified Eagle medium (KO-DMEM) supplemented with 20% KnockOut Serum replacement (Thermo Fisher Scientific), 1% non-essential amino acids (MEM-NEAA) (Thermo Fisher Scientific), 1% GlutaMAX (Thermo Fisher Scientific), 0.1% β-mercaptoethanol (Thermo Fisher Scientific) and 8ng/ml human recombinant basic fibroblast growth factor (Sigma Aldrich). Cultures were routinely tested for mycoplasma using a PCR assay^79^. All feeder-free experiments were performed from cultures grown on hESC-qualified Matrigel (Corning) and maintained in mTeSR plus media (STEMCELL Technologies).

### Immunofluorescence microscopy and analysis

hESCs were fixed in 4% paraformaldehyde (in PBS) for 5 minutes, permeabilized with 0.1% Triton-X 100 and blocked using 4% fetal bovine serum (FBS) for 1 hr at RT (room temperature). Cells were then incubated with the appropriate primary antibodies overnight at 4°C, washed with PBS and incubated with secondary antibody conjugated to either Alexa Fluor 488 or 568 for 1 hr at RT. The cells were stained using 10μg/ml DAPI (Sigma Aldrich) to visualize nuclei. Imaging was performed using either an inverted epifluorescence microscope (Olympus IX-81) equipped with an EM CCD camera (Andor Luca R) or a sCMOS camera (ORCA Flash 4.0 v3, Hamamatsu), or a Zeiss LSM880 confocal microscope.

The antibodies used in this study are listed in supplemental table 1. Images were processed uniformly for brightness and contrast using Adobe Photoshop CS5.

For nuclear SMAD2/3 and p-SMAD1/5/8 analysis, 3 or 6 hrs mesendoderm differentiating hESC colony edges were imaged to quantify signalling activation. Maximum intensity projections were generated using LSM software and exported as 8-bit files into ImageJ (NIH). The nuclear region was identified using DAPI staining, and mean fluorescence intensity was quantified specifically within the DAPI-marked nuclear area. For quantification of Pax6^+^ cells, 8-bit image projections were thresholded and quantified using the Analyze particle function in ImageJ (NIH). This was expressed as % Pax6^+^ cells, normalized to the total number of DAPI-stained nuclei.

### Live imaging and analysis of the mitochondrial network

Live tracking and imaging of the mitochondrial network was performed as done earlier^24^. Z-projected images were analysed using the Mitochondrial Network Analysis Macro tool (MiNA plugin) in Fiji, as described previously^80^, to quantify mitochondrial network parameters, including branch length, branch number, junction number and footprint. The mitochondrial footprint per cell was estimated by quantifying the total mitochondrial footprint in the region of interest and normalizing it to the number of nuclei identified in the corresponding phase-contrast images. The mitochondrial network dynamics was assessed by calculating the variance of these mitochondrial morphology parameters over time, as done previously^81^.

### RNA isolation and reverse transcription-quantitative PCR (RT-qPCR)

RNA extraction was performed using TRIzol reagent (Life Technologies) as per the manufacturer’s instructions. 4-6μg of DNase-treated total RNA was reverse transcribed to cDNA using the Superscript III first strand synthesis kit (Life Technologies) and random primers (Invitrogen). RT-qPCR was set up with either SsoFast EvaGreen Supermix, (Bio-Rad) or SensiFAST SYBR No-ROX Kit (Bioline). The primers used in this study are listed in supplemental table 2.

### Immunoblotting

Total protein of hESCs was extracted using a lysis buffer consisting of 20 mM HEPES (pH 7.5), 150 mM NaCl, 10% glycerol, 5 mM MgCl_2_, 5 mM EDTA (pH = 8), 0.5% Triton X-100, 1 mM PMSF, 1 mM sodium orthovanadate, 1X protease inhibitor cocktail (Sigma Aldrich) and 10μM proteasomal inhibitor MG-132 (Sigma Aldrich). The cells were incubated in the lysis buffer for 6 hrs at 4°C in an end-to-end rotor, followed by centrifugation at 12,000 rcf for 30 minutes at 4°C. The supernatant was collected and total protein was estimated using the Bradford reagent (Bio-Rad). 40μg of the denatured lysate was used for standard immunoblotting analysis. Antibodies used in this study are listed in Table 1. Signals were developed through appropriate horse-radish peroxidase (HRP)-conjugated secondary antibodies (Genei), using an enhanced chemiluminescence (ECL) kit (Bio-Rad) and visualized on X-ray films in the dark room. Band intensities were quantified using Fiji software and normalized to Vinculin or α-Tubulin as the loading control.

### Mesendoderm differentiation and small molecule treatments

Directed mesendoderm differentiation was performed as described previously^40^. Briefy, hESCs were mechanically passaged, seeded and allowed to attach on Matrigel coated dishes for 48 hrs. Upon reaching 50-60% confluency, the media was replaced by StemLine II hematopoietic stem cell expansion medium (Sigma Aldrich) supplemented with growth factors such as Activin A (Sigma Aldrich), BMP4 (Sigma Aldrich), VEGF (Sigma Aldrich) and 10ng/mL bFGF (Sigma Aldrich), each 10ng/mL. Activin A was included only for the first two days of induction, unless indicated otherwise. The differentiation media was refreshed every two days and cells were harvested for analysis as required. Small molecules such as SB431542 hydrate (5μM or 10μM) (Sigma Aldrich), 2μM Dorsomorphin (Sigma Aldrich) were used to inhibit TGFβ and BMP4 pathway respectively while 5μM CHIR99021 (Sigma Aldrich) was used for activating WNT pathway. For sodium acetate and etomoxir experiments, attached hESC aggregates were pre-treated with 10mM sodium acetate (Sigma Aldrich) or 100μM Etomoxir sodium salt hydrate (Sigma Aldrich) in mTeSR plus medium for 24 hrs, followed by mesendoderm induction media supplemented with 10mM sodium acetate or 100μM Etomoxir sodium salt hydrate until day 3.5. Directed mesoderm induction of hESCs was carried out by inducing hESC aggregates with STEMdiff^TM^ Mesoderm Induction Medium (STEMCELL technologies) for 4 days.

### Neuroectoderm differentiation

Neuroectoderm differentiation was performed following the described protocol^82^. Briefly, hESCs were seeded as a monolayer to reach 90% confluency to begin differentiation. The neuroectoderm induction contained DMEM/F-12 (Invitrogen), 450μM 1-thioglycerol (Sigma Aldrich), 0.1% Bovine Serum Albumin (BSA) (Sigma Aldrich), 0.11μM β-mercaptoethanol (Thermo Fisher Scientific), 1% B-27 supplement (Thermo Fisher Scientific), 1% N2 supplement (Thermo Fisher Scientific), 10μM SB431542 and 0.2μM Dorsomorphin. The media was replenished daily and neuroectoderm potential was evaluated on day 8 post-induction by Pax6 immunostaining.

### Flow cytometry analysis

BJNhem20 cells grown on Matrigel coated dishes were harvested using TrypLE Express enzyme (Life Technologies), washed with 1X PBS and taken ahead for staining. The fluorochrome-conjugated antibodies used in this study are listed in supplemental table 1. For staining with fluorescent dyes, cells were stained with 100μM 2-NBDG (Invitrogen) or 5μM BODIPY 505/515 (4,4-Difluoro-1,3,5,7-Tetramethyl-4-Bora-3a,4a-Diaza-s-Indacene) (Invitrogen) in 1X PBS for 30 min at 37°C. Samples were subjected to flow cytometry on a FACS Aria III (BD Biosciences) and 10,000 or 50,000 events were recorded in triplicates. Data obtained was analysed using FlowJo v10 software (BD).

### Ki-67 Proliferation and Annexin V apoptosis assay

BJNhem20 lines were harvested as mentioned previously, and assayed for Ki-67 staining using the FITC Ki-67 proliferation kit (BD Biosciences), according to the manufacturer’s instructions. Cells were co-stained with 20μg/mL Hoechst 33342 (Thermo Fisher Scientific) to mark the different cell cycle stages. Assessment of apoptosis was done using the FITC Annexin V Apoptosis Detection Kit I (BD Biosciences), following the manufacturer’s protocol. Propidium idodide was used to mark dead cells. Samples were analysed using flow cytometry.

### Wound healing assay

hESCs were seeded at 50,000/cm^2^ in a 24-well multiplate and allowed to grow for 36 hrs to form a monolayer. Cells were then induced to differentiate for 6 hrs using mesendoderm media. After 6 hrs of induction, cells were treated with 10μg/mL mitomycin C for 2 hrs to block proliferation. The monolayer was scratched down the center using a sterile 20μL pipette tip and washed with medium to remove traces of mitomycin C and debris. Scratches were imaged directly in the center of the colony at time 0 hr and 18 hrs post-scratching by phase-contrast microscopy. The images were analysed in ImageJ (NIH) using the wound healing tool plugin (RRID:SCR_025260). The extent of migration (closure) was quantified by calculating percent change in the scratched area covered by cells in 18 hrs.

### Bulk RNA sequencing and analysis

Total RNA was extracted from either undifferentiated or day 2 mesendoderm-induced hESCs by the TriZol method. RNA was quantified using a Qubit 4 fluorimeter (Invitrogen) and its quality was checked with a TapeStation system (Agilent) using the high-sensitivity RNA Screen Tape analysis (Agilent Technologies). mRNA enrichment was performed with Next Poly(A) mRNA Magnetic Isolation Module (New England Biolabs). cDNA library was prepared using the Next® Ultra™ II Directional RNA Library Prep with Sample Purification Beads (New England Biolabs). The libraries were sequenced on the NovaSeq 6000 platform (Illumina) with 2×100 bp sequencing read length, generating over 50 million paired-end reads per sample.

Raw fastq files were checked for their data quality using FastQC tool^83^, both before and after adapter trimming, and were finally aligned to the human reference genome (GRCh38) with HISAT2^84^. Gene expression counts were determined by the Feature Counts program of the Subread R package, using the default parameters^85^. Differential expression analyses were performed with the R package DESeq2 (v1.30.0)^86^. Significance of differentially expressed genes was defined by setting the threshold: adjusted p-value < 0.05 and abs | log_2_(fold change) | > 0.5.

Principal Component Analysis (PCA) was performed using the prcomp function in R. Volcano plots were generated using the ggplot2 package in R and heatmaps were constructed using pheatmap or complex heatmap packages. Gene ontology (GO) analysis was performed using the clusterProfiler package (v4.12.6)^87^ and barplots were constructed in Prism 8.3 (GraphPad) for Windows. Gene Set Enrichment Analysis (GSEA) was done with the GSEA tool (4.3.3) using MSigDB database^88^. Pathway enrichment analysis was carried out using Kyoto Encyclopedia of Genes and Genomes (KEGG) database in the Enrichr web tool^89^.

### Seahorse extracellular flux assays

5,000 hESCs were allowed to attach and grow on Matrigel coated 24-well FluxPak mini pack plates (Agilent Technologies) for 72 hrs at 37°C with 5% CO2. The oxygen consumption rate (OCR) was computed using the Seahorse XF Mito stress Test kit (Agilent Technologies). The assay medium consisted of unbuffered DMEM (Agilent Technologies) supplemented with 1% GlutaMAX (Thermo Fisher Scientific). The mito stress assay was done using 1μM oligomycin, 0.3μM FCCP (carbonyl cyanide p-trifluoromethoxy phenylhydrazone), and 1μM each of rotenone and antimycin A (Rot + AA.). Flux analysis was done on Seahorse bioanalyzer XFe24 with three-minute measurement cycles. The OCR values were normalized to relative viable cells present in each well, quantified using Prestoblue (Thermo Fisher Scientific). OCR values were averaged over technical replicates and represented from three independent experiments.

### Statistical analysis and data representation

All data are presented as mean ± standard error of the mean (SEM) calculated from at least three independent experiments. Statistical significance was determined using either one sample two-tailed t-test, ratio paired t-test, one-way ANOVA with Dunette’s or two-way ANOVA with Tukey’s multiple comparisons test. For samples with unequal variance, the Kruskal Wallis non-parametric test was performed. All statistical analyses and graph plotting were done in Prism 8.3 software for Windows (GraphPad).

## Data and code availability

All data supporting the findings of this study are included in the main text and supplemental figures. Bulk RNA-seq datasets can be accessed via GEO: GSE307502. The original raw data and analysis files are available from the corresponding authors upon reasonable request.

## Acknowledgements

We thank the biology research facility at JNCASR, next generation genomics facility at National Centre for Biological Research (NCBS), and central facilities at iBRIC-inStem. This work was funded by the Department of Biotechnology, Government of India (grant no. BT/PR23947/MED/31/372/2017), JC Bose Fellowship (JCB/2019/000020) to MI and intramural funds from Jawaharlal Nehru Centre for Advanced Scientific Research (JNCASR), Bangalore and Institute for Stem Cell Science and Regenerative Medicine (iBRIC-inStem). Schematics were generated using Biorender.

## Author contributions

Conceptualization: KK and MI; data curation, formal analysis, validation, investigation, visualization, methodology: KK; writing - original draft, review and editing: KK and MI; funding acquisition, resources and supervision: MI.

## Declaration of interests

The authors declare no competing interests.

---

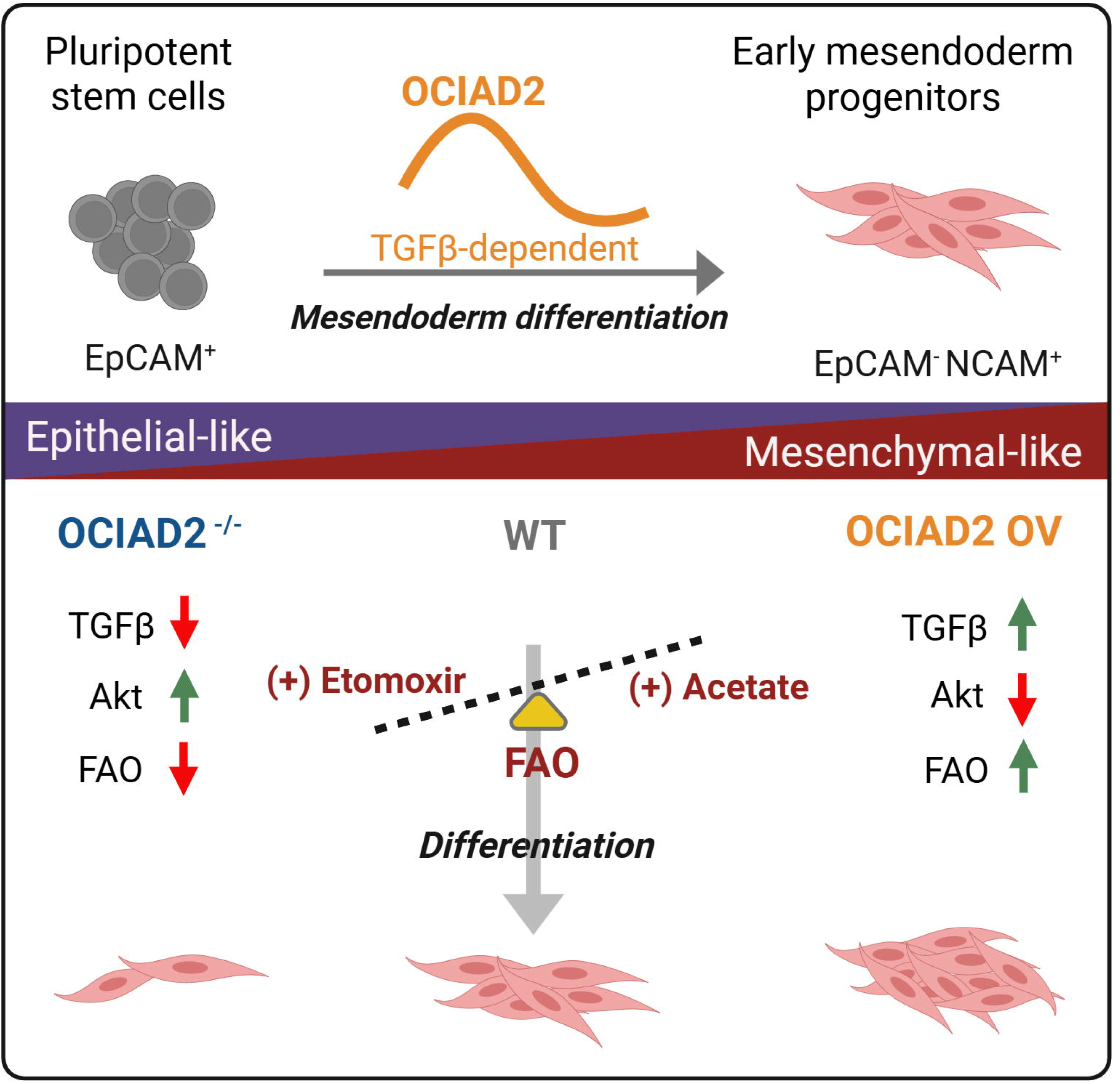

## Notes

### Competing Interest Statement

The authors have declared no competing interest.

