## Supplemental information for "TGFβ-dependent upregulation of OCIAD2 is essential for epithelial-to-mesenchymal transition during mesendoderm differentiation"

Supplemental text, figures and tables: Supplemental figures S1-S6, supplemental figure legends, and supplemental tables S1-S2.

Supplemental video S1.

**Supplemental figure legends**

**Figure S1. Undifferentiated hESCs express OCIAD2 (*Related to figure 1*)*.***

**(A)** Immunostaining analysis of OCIAD2 with pluripotency markers Oct3/4, TRA-1-60, TRA1-81. Nucleus is marked by DAPI. Scale bar: 50μm.

**(B)** RT-PCR analysis showing *OCIAD2* transcript expression in hESCs.

**(C)** Immunoblotting of OCIAD2 in undifferentiated hESCs.

**(D)** UMAP projection of *OCIAD2* expression in integrated human embryonic datasets representing developmental stages from the zygote to the CS7 gastrula^1^. Source for UMAP visualization- <https://petropoulos-lanner-labs.clintec.ki.se/shinys/app/ShinyEmbryoRef>

**(E)** Schematic representing *OCIAD2* expression obtained from bulk transcriptome of directed mesoderm differentiated hESCs^2, 3^. Each box denotes a cell-type generated during human mesoderm development and the intensity of shading represents the expression in that cell-type. Scale indicates log_2_ gene expression. Schematic generated using the web tool- <https://z-chen.net/tree>

**Figure S2. OCIAD2 levels remain unaffected upon inhibition of BMP signalling (*Related to figure 1)*.**

**(A)** Schematic of the adapted mesendoderm induction protocol^4^, indicating the treatment windows for targeting Activin A and BMP signalling.

**(B)** *OCIAD2* transcript expression across days of mesendoderm specification.

Increase in *NCAM1* levels mark onset of mesendoderm specification. Transcript levels were normalized to *GAPDH*. Graph represents fold change in transcript expression.

**(C)** RT-qPCR analysis of *OCIAD2*, *NANOG* and *CDH1* transcripts in undifferentiated hESCs upon TGFβ pathway inhibition. hESCs were treated with 10μM SB431542 (SB), a TGFβ type I receptor blocker for 48 hrs. Graph represents fold change in transcript expression normalized to RPLP0.

**(D)** Immunostaining of phospho-SMAD1/5/8 (p-SMAD1/5/8) in hESCs after 3 hrs of mesendoderm induction to confirm BMP pathway inhibition by 2μM Dorsomorphin (DM). Scale bar: 200μm.

**(E-F)** Immunoblotting for OCIAD2 levels upon blocking BMP4 pathway using 2μM Dorsomorphin (DM) from **(E)** day 0 to day 2.5 or **(F)** day 2 to day 4.5. Graphs show quantification of relative OCIAD2 levels normalized to vinculin.

Results shown are a representative of three independent experiments. Statistical significance was calculated using one sample two-tailed t-test (for **B**-**C**) or ratio paired two-tailed t-test (for **E-F**). Error bars denote standard error of mean, **p*<0.05, ***p*<0.01, ns indicates non-significant.

**Figure S3. OCIAD2 modulation does not affect stemness, proliferation and apoptosis in hESCs (*Related to figure 2*).**

**(A)** Flow cytometry analysis of the undifferentiated marker SSEA4 in OCIAD2 modulated hESCs. Histogram with the dotted outline represents the isotype control. Graph shows the quantification of % SSEA4^+^ cells.

**(B)** Flow cytometry analysis of Ki-67/Hoechst 33342 in OCIAD2 modulated hESCs. Boxed regions denote the different cell cycle stages identified using Hoechst 33342. Bar graph represents the frequency of these cell cycle stages.

**(C)** Flow cytometry analysis of Annexin V/propidium iodide (PI) staining. Annexin V⁺ PI⁻ cells represent early apoptotic cells, while Annexin V⁺ PI⁺ cells indicate late apoptotic cells.

**(D)** Gene Ontology (GO) analysis of positively enriched biological processes in day 0 OCIAD2 KO hESCs.

**(E)** Bar graph showing top enriched pathways in OCIAD2 KO hESCs identified using KEGG pathway enrichment analysis.

**(F)** Gene set enrichment analysis (GSEA) plot for WNT/β-catenin and TGFβ pathway genes in day 0 WT and OCIAD2 OV cells. NES value indicates pathway enrichment. p-value = 0.0 indicates statistically significant.

Results shown are a representative of three independent experiments. Statistical analysis was performed using One-way ANOVA with Dunette’s multiple comparisons test. Error bars denote standard error of mean, ns indicates non-significant.

**Figure S4. Effect of OCIAD2 modulation on mesoderm and neuroectoderm differentiation potential of hESCs (*Related to figure 3)*.**

**(A)** Graph showing quantification of % EpCAM^-^ NCAM^+^ cells (EMPs) at day 3.5 in mesendoderm induced WT and OCIAD2 KO hESCs treated with 5μM CHIR99021. DMSO was used as a vehicle control.

**(B)** Graph representing quantification of % EpCAM^-^ NCAM^+^ cells (EMPs) at day 4 in mesoderm induced cultures using STEMdiff^TM^ media.

**(C)** Representative confocal images of Pax6 expression at day 8 post-neuroectoderm differentiation. Scale bar: 100μm. Graph shows quantification of percentage of Pax6^+^ cells, normalized to DAPI stained cells.

Results shown are a representative of three independent experiments. Statistical analysis was performed using two-way ANOVA with Tukey’s multiple comparisons test (for **A-B**) or one-way ANOVA with Dunette’s multiple comparisons test (for **C**). Error bars denote standard error of mean, ****p*<0.001, *****p*<0.0001, ns indicates non-significant.

**Figure S5. OCIAD2 modulates TGFβ and PI3K-AKT signalling in hESCs (*Related to figure 4*).**

**(A)** Heatmap analysis of mesendoderm-induced day 2 WT, OCIAD2 KO and OCIAD2 OV RNA-seq data representing all significant DEGs [log_2_ FC > ± 0.5; p_adj_ value < 0.05].

**(B)** Bar graph showing top enriched pathways in OCIAD2 OV hESCs identified using KEGG pathway enrichment analysis.

**(C)** Heatmap depicting the expression of TGFβ pathway genes in undifferentiated OCIAD2 modulated hESCs.

**(D-E)** Immunoblotting of **(D)** total SMAD2/3 and phospho-SMAD2 (Serine 465/467) and **(E)** total AKT and phospho-AKT (Serine 473) levels in undifferentiated OCIAD2 modulated hESCs. Graphs show quantification of fold change in protein levels normalized to vinculin.

Results shown are a representative of three independent experiments. One sample two-tailed t-test was used for statistical comparison. Error bars denote standard error of mean, **p*<0.05, ***p*<0.01, ns indicates non-significant.

**Figure S6. OCIAD2 regulates mitochondrial dynamics but does not affect OXPHOS in hESCs (*Related to figure 5*).**

**(A)** Violin plots represent variance in mitochondrial footprint, branch number and junction number across time, indicating dynamicity of the network in OCIAD2 modulated hESCs.

**(B)** Oxygen consumption rate (OCR) profile of OCIAD2 modulated hESCs measured using Seahorse Mito stress assay. Graph shows maximal respiration (induced by FCCP) and ATP-linked respiration (inhibited by oligomycin). Each data point represents the mean OCR value from an independent experiment.

**(C)** Histogram overlay representing 2-NBDG uptake in day 0 OCIAD2 modulated hESCs. Histogram with the dotted outline represents the FMO control. Graph shows quantification of median fluorescence intensity (MFI) of 2-NBDG.

**(D-E)** Graphs showing quantification of % EpCAM^+^ cells, %EMPs and NCAM MFI at day 3.5 in **(D)** WT and/or OCIAD2 KO hESCs treated with 10mM sodium acetate and **(E)** WT and/or OCIAD2 OV treated with 100μM etomoxir.

Results shown are a representative of three independent experiments. Statistical comparisons were performed using Kruskal Wallis test (for **A**), One-way ANOVA with Dunette’s multiple comparisons test (for **B**), One sample two-tailed t-test (for **C**) and One-way ANOVA with Dunette’s multiple comparisons test (for **D, E)**. Error bars denote standard error of mean, **p*<0.05, ***p*<0.01, *****p*<0.0001, ns indicates non-significant.

**Supplemental tables**

**Table S1: List of antibodies used in this study**

| **Sr.No.** | **Antibody** | **Company, Cat. No.** | **Application** |
| --- | --- | --- | --- |
| 1. | E-cadherin | BD Biosciences, Cat. no. 610182 | WB |
| 2. | N-cadherin | BD Biosciences, 610921 | IF |
| 3. | OCIAD2 | Sigma Aldrich, HPA041090 | WB |
| 4. | OCT3/4 | BD Biosciences, 611203 | IF |
| 5. | Pan-Akt | Cell Signalling Technology, 4691S | WB |
| 6. | p-Akt | Cell Signalling Technology, 4060S | WB |
| 7. | p-SMAD2 | Cell Signalling Technologies, 18338S | WB |
| 8. | p-SMAD1/5/8 | Cell Signalling Technology, 9511 | IF |
| 9. | SMAD2/3 | Cell Signalling Technology, 8685S | WB/IF |
| 10. | TRA1-60 | Kind gift from Prof. Peter Andrew, University of Sheffield, UK | IF |
| 11. | TRA1-81 | Kind gift from Prof. Peter Andrew, University of Sheffield, UK | IF |
| 12. | Vimentin | Sigma Aldrich, V6630 | IF |
| 13. | Vinculin | Sigma Aldrich, V4505 | WB |
| 14. | α-Tubulin | Sigma Aldrich, T8203 | WB |
| 15. | SSEA4-PerCP-Cy5.5 | BD Biosciences, 561565 | Flow cytometry |
| 16. | Isotype IgG3-PerCP-Cy5.5 | BD Biosciences, 561572 | Flow cytometry |
| 17. | NCAM (CD56)-PE | Biolegend, 318306 | Flow cytometry |
| 18. | Isotype IgG1-PE | Biolegend, 400114 | Flow cytometry |
| 19. | EpCAM  (CD326)- PerCP-Cy5.5 | Biolegend, 324214 | Flow cytometry |
| 20. | Isotype IgG2b-PerCP-Cy5.5 | Biolegend, 400338 | Flow cytometry |
| 21. | FITC Ki-67 proliferation kit, Ki-67-FITC antibody | BD Biosciences, 556026 | Flow cytometry |
| 22. | FITC Annexin V  Apoptosis Detection  Kit I, Annexin V-FITC  antibody | BD Biosciences, 556547 | Flow cytometry |

**Table S2: List of primer sequences used in this study**

| **Sr. No.** | **Primer name** | **Sequence: 5’ to 3’** |
| --- | --- | --- |
| 1. | *OCIAD2_F*  *OCIAD2_R* | TTCAGCGTCTGCTCGTGG  AACCTTGGTAGACTAGTCCC |
| 2. | *NCAM1_F*  *NCAM1_R* | GCCAGGAGACAGAAACGAAG  GGTGTTGGAAATGCTCTGGT |
| 3. | *CDH1_F*  *CDH1_R* | TGCCCAGAAAATGAAAAACG GTGTATGTGGCAATGCGTTC |
| 4. | *MIXL1_F*  *MIXL1_R* | TCCAGGATCCAGGTATGGTT  GCTCCTCAGAGCTTATCCCGAA |
| 5. | *GAPDH_F*  *GAPDH_R* | CCCATGTTCGTCATGGGTG  GATGGCATGGACTGTGGTC |
| 6. | *RPLP0_F*  *RPLP0_R* | CACCATTGAAATCCTGAGTGATGT  CTGCCATTGTCGAACACCTGCT |
| 7. | *SNAI2_F*  *SNAI2_R* | GGGGAGAAGCCTTTTTCTTG  TCCTCATGTTTGTGCAGGAG |
| 8. | *VIM_F*  *VIM_R* | GAGAACTTTGCCGTTGAAGC  GCTTCCTGTAGGTGGCAATC |
| 9. | *EOMES_F*  *EOMES_R* | AAATGGGTGACCTGTGGCAAAGC  TTGTGTAAGGATTGTAAGACTAT |
| 10. | *GSC_F*  *GSC_R* | GAGGAGAAAGTGGAGGTCTGG GCAAGAAAGTAGCATCGTCTG |
| 11. | *NANOG_F*  *NANOG _R* | TGCAAATGTCTTCTGCTGAGAT  GTTCAGGATGTTGGAGAGTTC |

**Supplemental video**

**Video S1. 3D projected images of mitotracker deep red labelled mitochondria in live OCIAD2 modulated hESCs with 3fps rotation *(related to figure 5, figure S6)*.**
