## Supplementary figures and images for "TGFβ-dependent upregulation of OCIAD2 is essential for epithelial-to-mesenchymal transition during mesendoderm differentiation"

### Figure S1

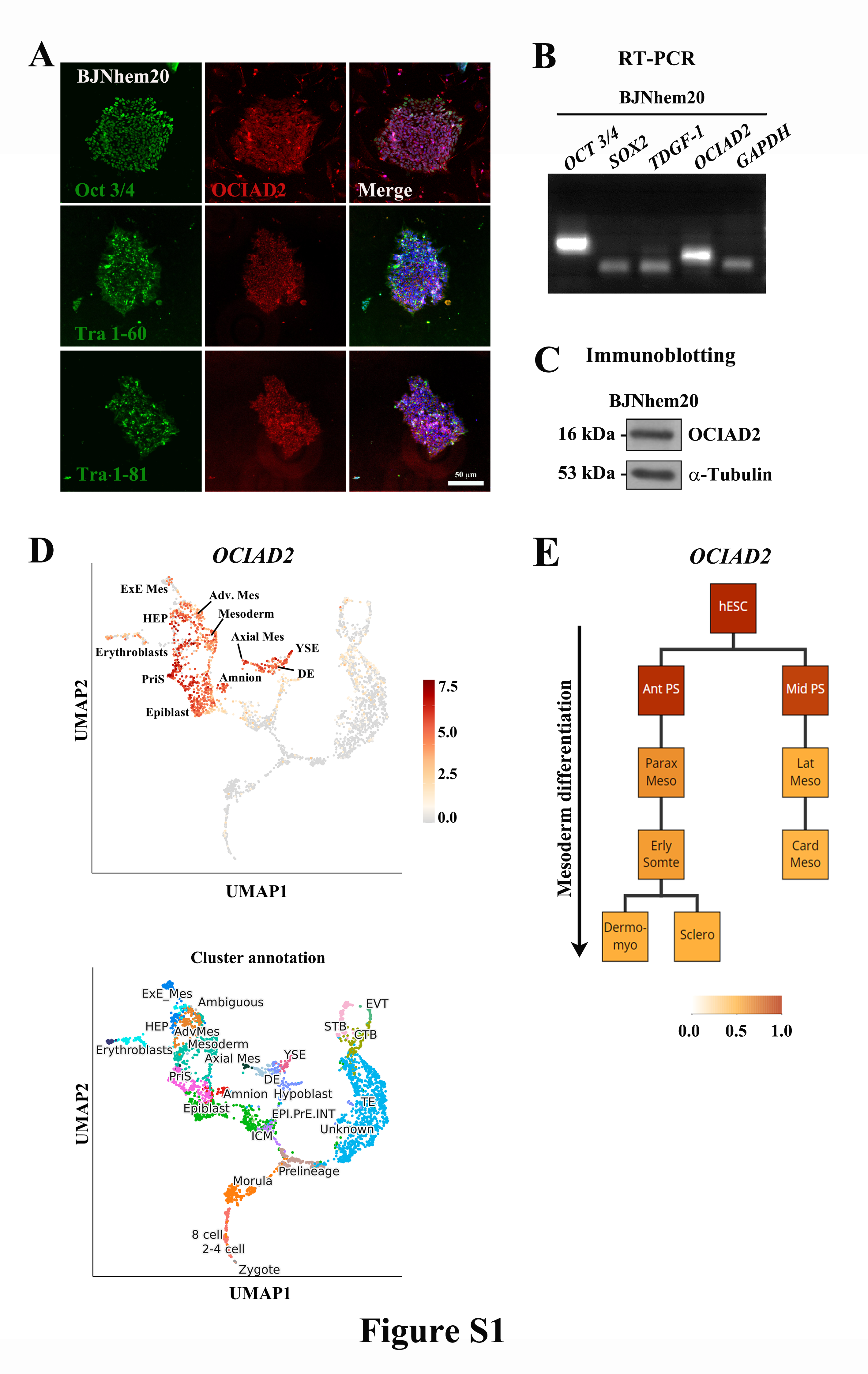

### Figure S2

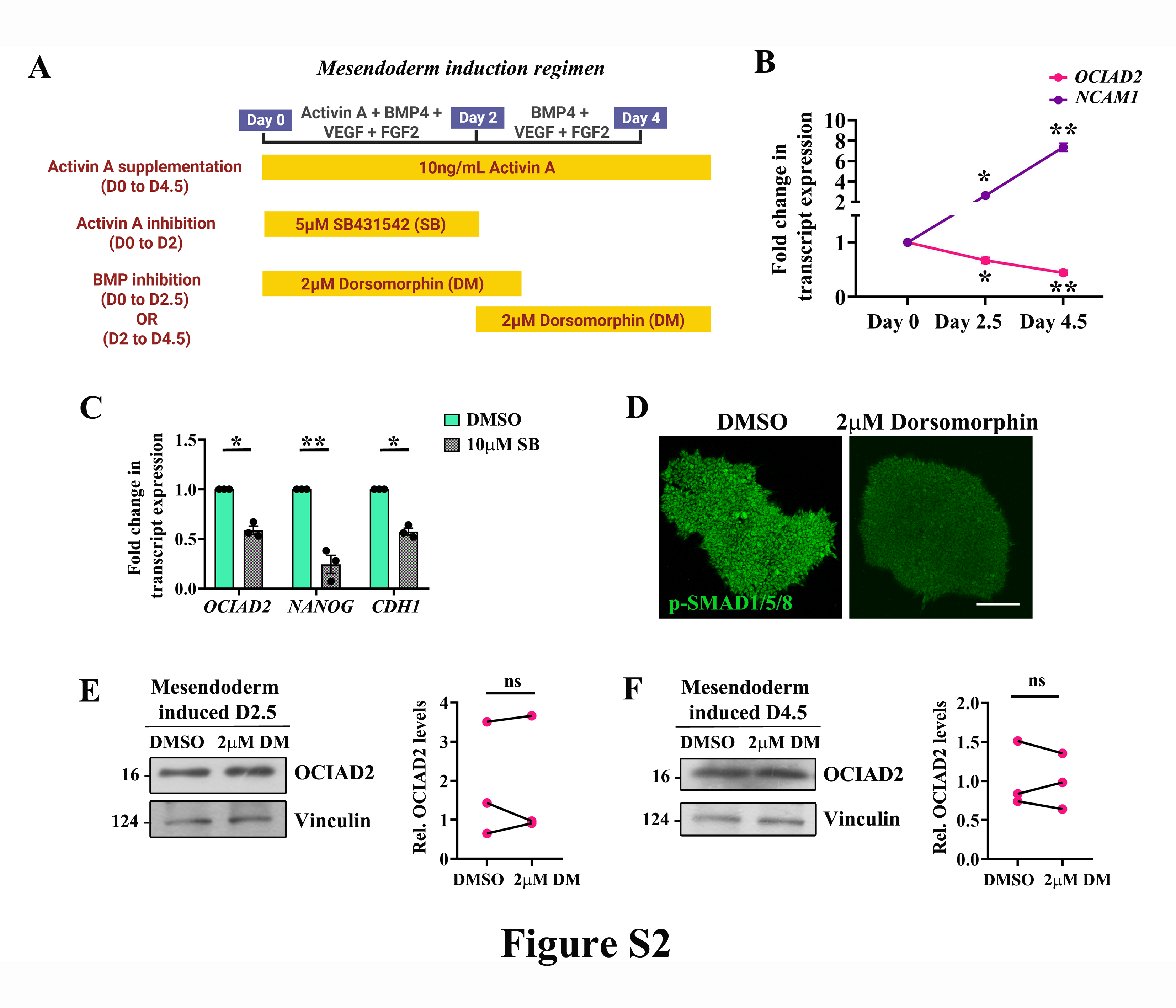

### Figure S3

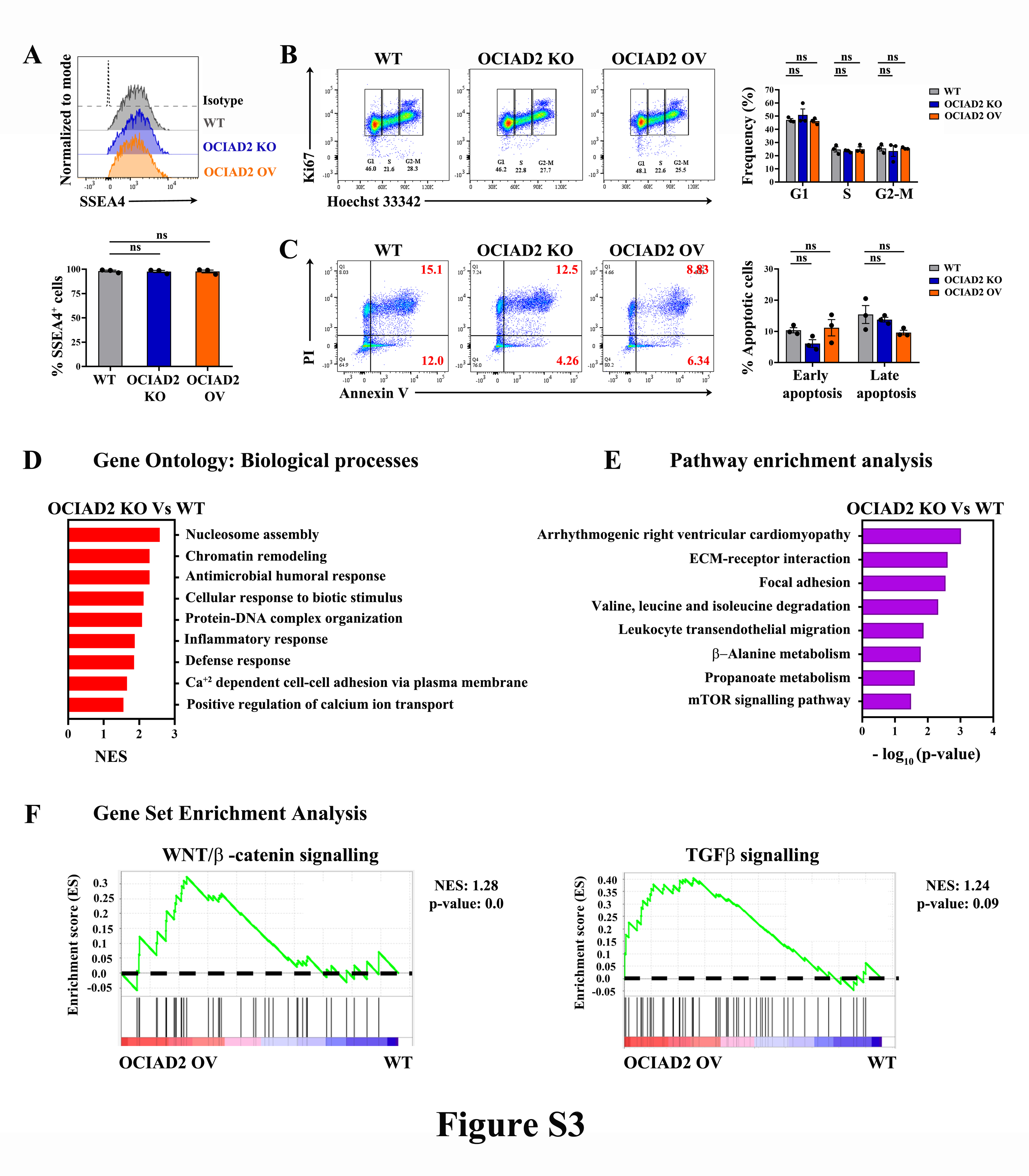

### Figure S4

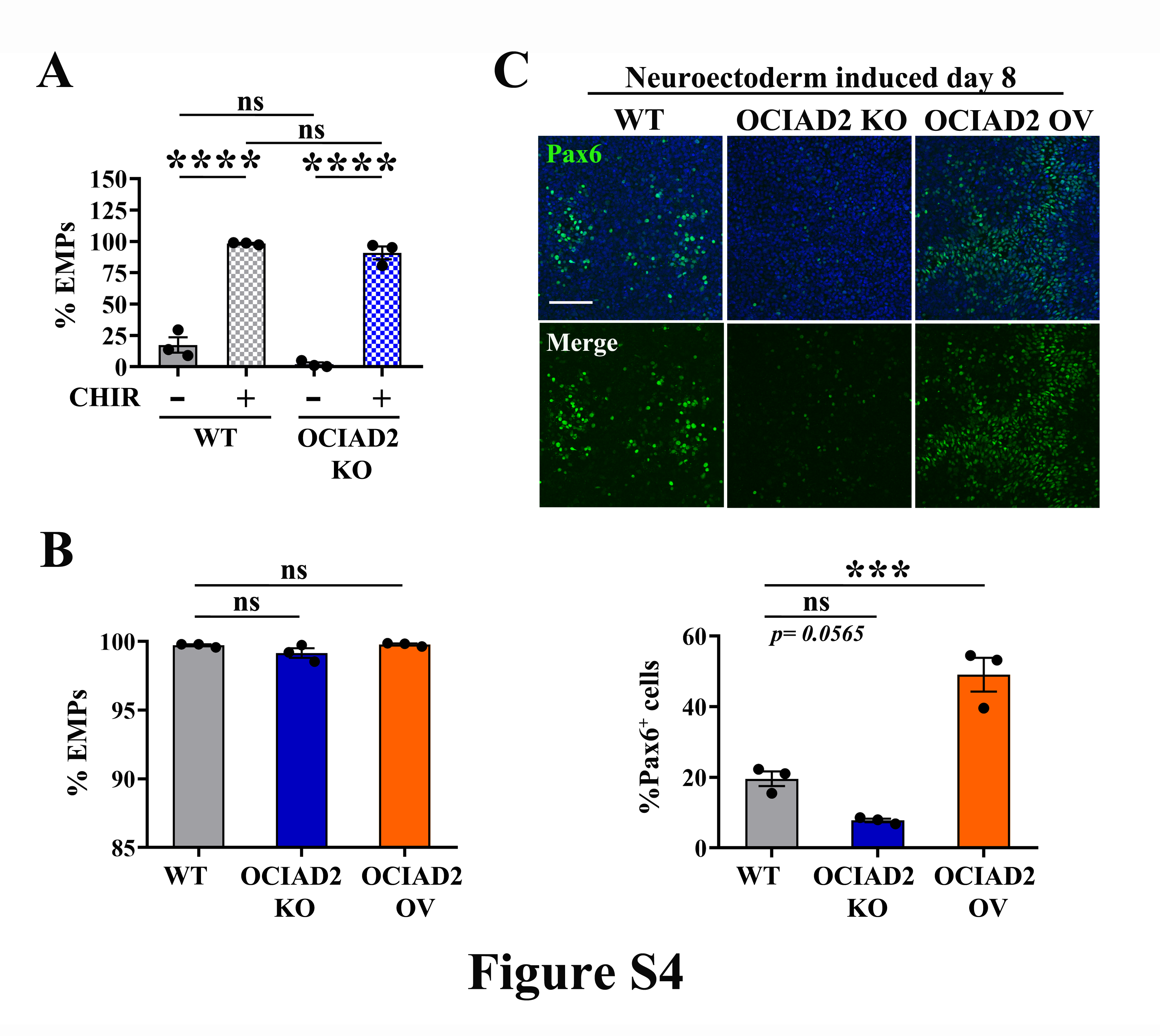

### Figure S5

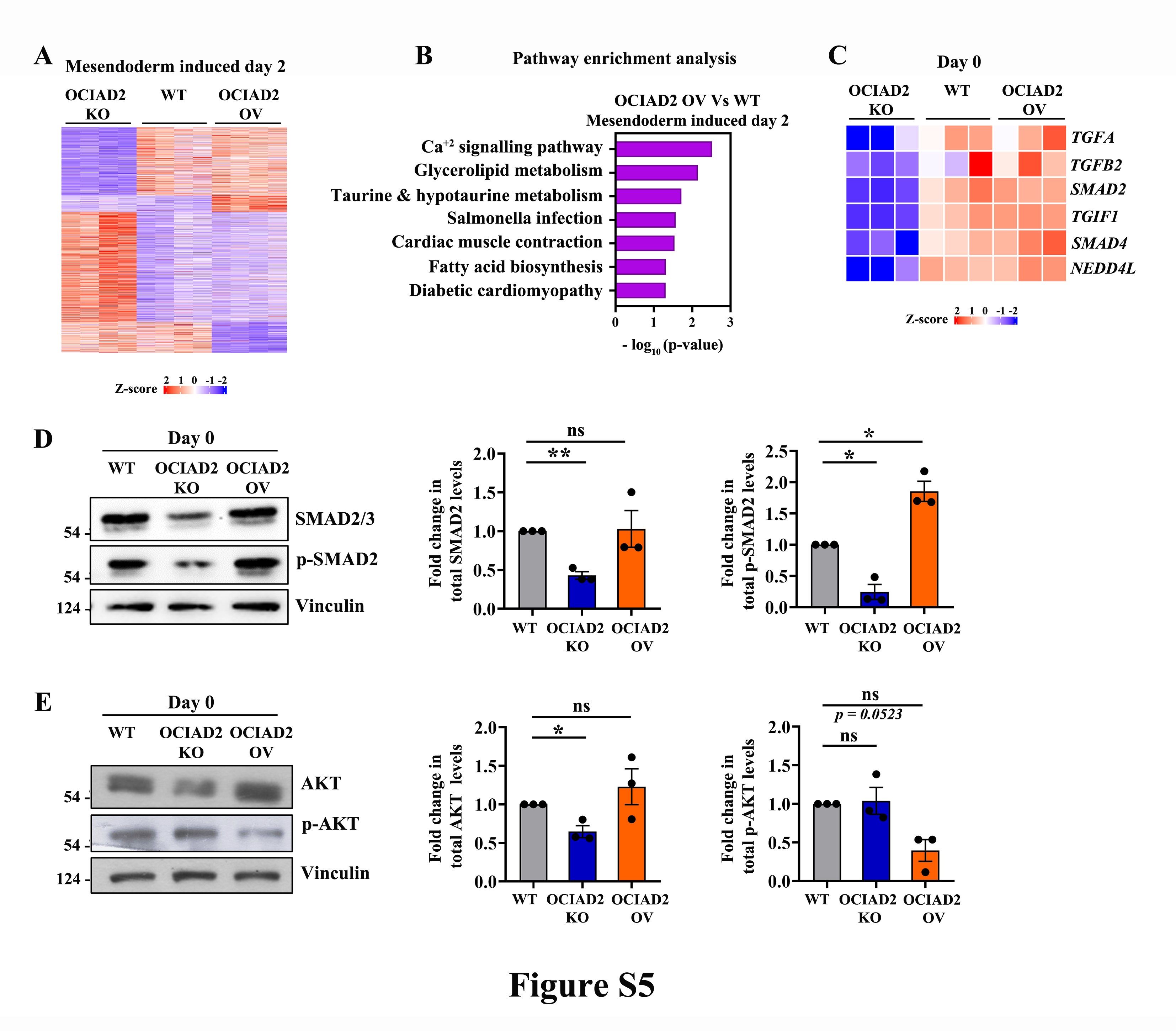

### Figure S6

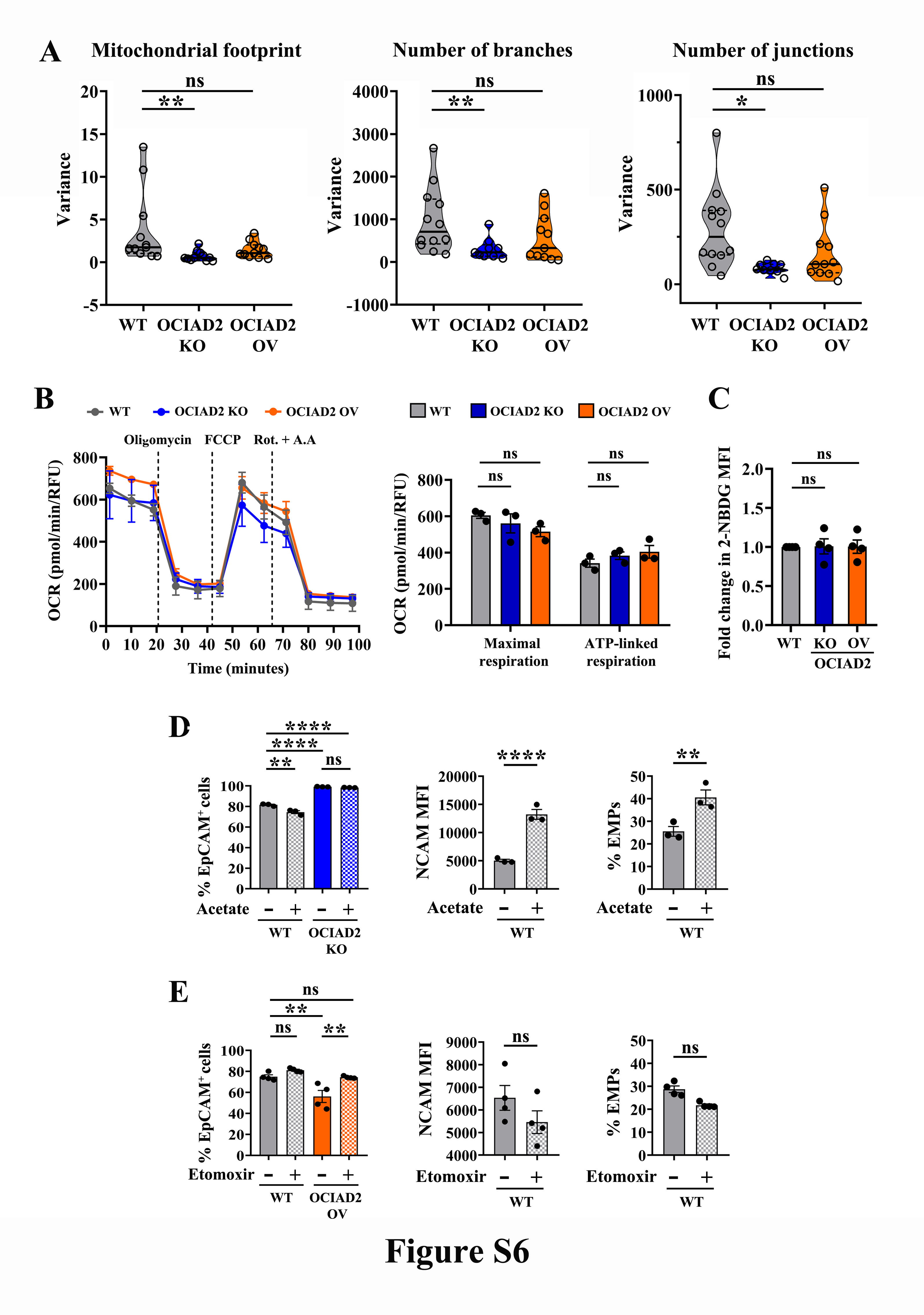
